# Genome-scale spatial mapping of the Hodgkin lymphoma microenvironment identifies tumor cell survival factors

**DOI:** 10.1101/2025.01.24.631210

**Authors:** Vignesh Shanmugam, Neriman Tokcan, Daniel Chafamo, Sean Sullivan, Mehdi Borji, Haley Martin, Gail Newton, Naeem Nadaf, Saoirse Hanbury, Irving Barrera, Dylan Cable, Jackson Weir, Orr Ashenberg, Geraldine Pinkus, Scott Rodig, Caroline Uhler, Evan Macosko, Margaret Shipp, Abner Louissaint, Fei Chen, Todd Golub

## Abstract

A key challenge in cancer research is to identify the secreted factors that contribute to tumor cell survival. Nowhere is this more evident than in Hodgkin lymphoma, where malignant Hodgkin Reed Sternberg (HRS) cells comprise only 1-5% of the tumor mass, the remainder being infiltrating immune cells that presumably are required for the survival of the HRS cells. Until now, there has been no way to characterize the complex Hodgkin lymphoma tumor microenvironment at genome scale. Here, we performed genome-wide transcriptional profiling with spatial and single-cell resolution. We show that the neighborhood surrounding HRS cells forms a distinct niche involving 31 immune and stromal cell types and is enriched in CD4+ T cells, myeloid and follicular dendritic cells, while being depleted of plasma cells. Moreover, we used machine learning to nominate ligand-receptor pairs enriched in the HRS cell niche. Specifically, we identified IL13 as a candidate survival factor. In support of this hypothesis, recombinant IL13 augmented the proliferation of HRS cells *in vitro*. In addition, genome-wide CRISPR/Cas9 loss-of-function studies across more than 1,000 human cancer cell lines showed that IL4R and IL13RA1, the heterodimeric partners that constitute the IL13 receptor, were uniquely required for the survival of HRS cells. Moreover, monoclonal antibodies targeting either IL4R or IL13R phenocopied the genetic loss of function studies. IL13-targeting antibodies are already FDA-approved for atopic dermatitis, suggesting that clinical trials testing such agents should be explored in patients with Hodgkin lymphoma.

## Introduction

Sustained growth factor signaling is a fundamental hallmark of cancer^1,2^. Most prior studies have focused on the somatic cancer genome as a source of insight into the mechanisms explaining such aberrant signaling, as similarly comprehensive analyses of the tumor microenvironment have been technically infeasible. Yet not all cancers have somatic mutations that readily explain tumor survival, and the fact that human tumors often contain high proportions of non-malignant cells (fibroblasts, lymphoid cells, and myeloid cells) suggests that such microenvironmental cells may contribute to tumor cell survival. Consistent with this hypothesis, establishing cancer cell lines is often difficult, if not impossible, presumably because the tumor cells cannot survive when deprived of microenvironment-derived survival factors.

Survival signals are particularly important in B-cell lymphomas because normal B cells are highly dependent upon signals transduced through cytokine and antigen receptors within lymph node follicles^3^. Malignant B cells can overcome the limiting availability of these growth signals through somatic mutations in the B cell receptor or its downstream effectors^4,5^, and therapeutics targeting this pathway have proven effective in clinical trials of diffuse large B-cell lymphoma (DLBCL)^6–8^.

Whereas DLBCL typically grows as monotonous sheets of malignant B cells, other B-cell lymphomas are composed of a mixture of malignant B cells and immune cells^9–11^, suggesting that a cell non-autonomous survival mechanism might be at play. The most extreme example of a microenvironment-associated cancer is classic Hodgkin lymphoma (cHL), where the rare malignant Hodgkin Reed Sternberg (HRS) cells that comprise only 1-5% of the tumor’s cellularity are embedded in an extensive inflammatory infiltrate^12^. Several lines of evidence suggest the importance of the Hodgkin lymphoma microenvironment for HRS cell survival (as opposed to simply being a reactive infiltrate). First, HRS cells are notoriously difficult to propagate *in vitro* or in immunodeficient mice, with the success rate in establishing HRS cell lines being <1%^13–15^. Second, when cHL tumors spread to non-lymphoid organs, the HRS cell microenvironment is reconstituted at those distant sites^10,11^. Third, somatic activating mutations that lead to constitutive activation of oncogenes are relatively uncommon in cHL^16–19^, suggesting that growth and survival signals may be derived from the HRS cell niche.

Importantly, HRS cells have been shown to have immune evasion mechanisms, including loss of function mutations in MHC class I and increased expression of PD-L1 through copy number gains^20,21^. Recent immunophenotyping studies using candidate markers have demonstrated that distinct subsets of PD-L1+ macrophages and CD4+ T cells (including regulatory CTLA-4+ and LAG3+ subsets) surround HRS cells^22–24^. Single-cell transcriptomic studies of cHL have been reported^24–26^, but those studies lacked the ability to capture certain myeloid cells (which can make up > 50% of tumor cellularity^27^) and stromal cells, and they lacked spatial resolution. Therefore, an unbiased, comprehensive analysis of the cHL microenvironment has been desperately needed by the field.

In this study, we use spatially resolved expression profiling to systematically define the molecular and cellular architecture in the intact cHL tumor microenvironment. We identify novel immune evasive and trophic signaling mechanisms and functionally validate a critical role for IL13 in promoting HRS cell survival, suggesting that targeting IL13 signaling may have therapeutic benefit for patients with Hodgkin lymphoma.

## Results

### Transcriptional programs and topology of HRS cells

We first generated a comprehensive single nucleus and spatially resolved RNA sequencing dataset (fig. 1a, supplementary table 1) using samples from newly diagnosed cHL patients (n = 12), including EBV-positive and EBV-negative cases, as well as from control reactive lymphoid nodes (RLN, n = 7). Histopathologic analysis, single-nucleus RNA sequencing (snRNAseq), and spatial profiling using slide-seqV2 were performed on serial sections from each specimen (≥ 2 replicates for each sample and each technology platform). Following quality control (see Methods, extended data fig. 1a-c, 2a, supplementary table 2, 3), the resulting dataset yielded 1.9 million spatially resolved RNA profiles and 324,855 single nucleus RNA profiles (fig. 1b, c).

**Fig. 1:**
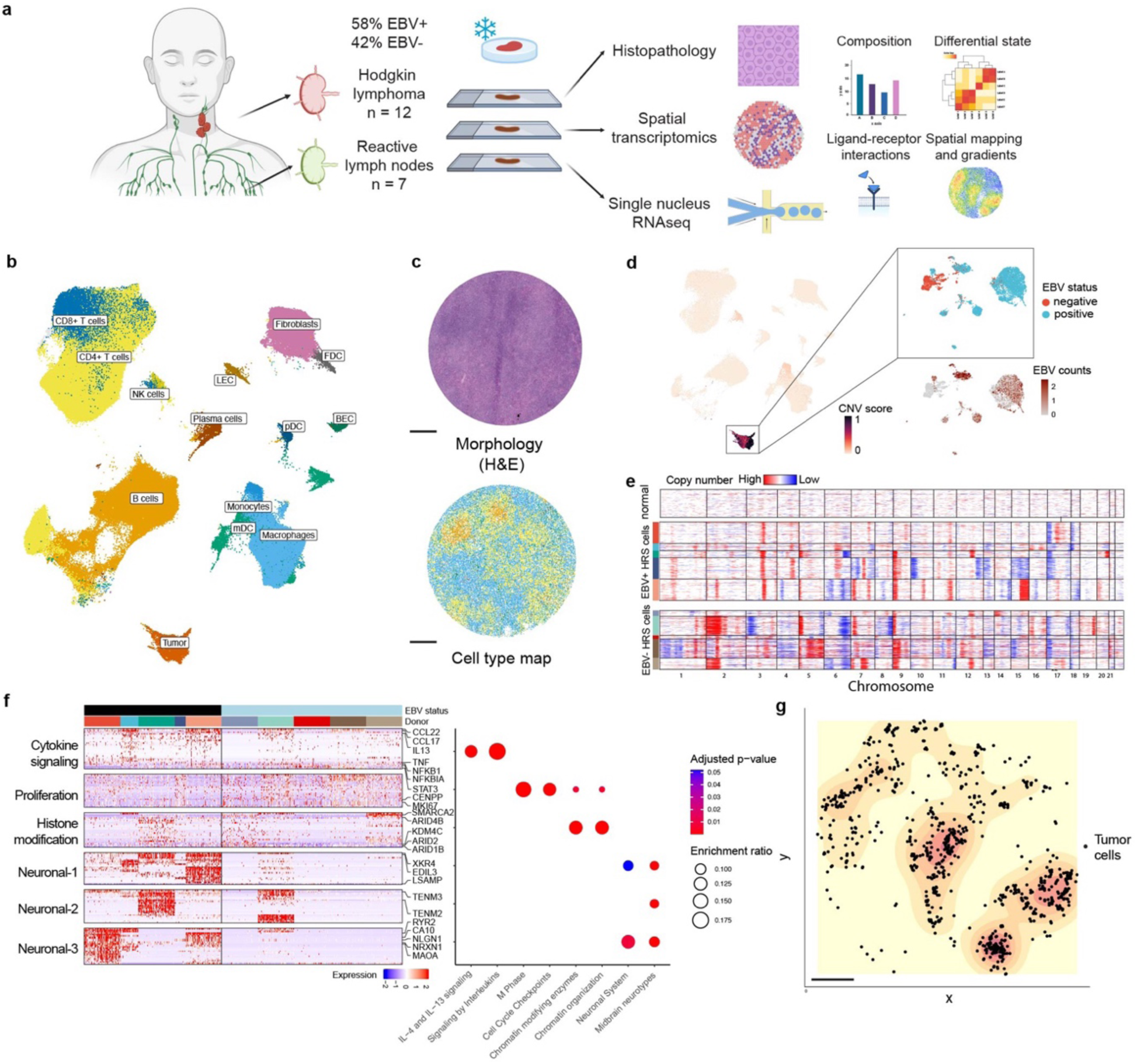
Transcriptional programs and topology of HRS cells. **a**, Schematic of experimental design and analyses. **b**, an overview of the single nucleus RNA sequencing (snRNAseq) dataset. Uniform manifold approximation and projection (UMAP) embedding of snRNAseq profiles colored by cell type annotation. **c**, representative slide-seqV2 array of a Hodgkin lymphoma tumor with hematoxylin and eosin (H&E) stained section (top) and spatial plot of cell types colored by cell type (bottom), same color scheme as b. **d**, identification of HRS cells by copy number variant (CNV) inference using inferCNV and EBV-specific read mapping. UMAP embedding showing CNV burden (quantified by CNV score) across cell types. The top right inset shows the UMAP embedding of HRS cells identified by high CNV burden colored by EBV status determined by in situ hybridization (positive or negative). The lower right inset shows the same UMAP embedding with the expression levels of EBV-specific transcripts. **e**, normalized and scaled gene expression arranged by chromosome number showing CNVs in EBV- and EBV+ HRS cells (grouped by donor) relative to non-tumor cells (top) that show no significant evidence of CNVs. **f**, Left: heatmap showing shared gene expression programs (normalized and scaled expression) in tumor cells across multiple donors identified by non-negative matrix factorization grouped by EBV status and donors. Right: enrichment analysis results of curated gene sets (columns) in each program (rows). **g**, spatial contour plot showing the clustered topology of HRS cells in tissue (representative array from donor H12-200). For c and g, scale bars, 500 μm.

We assigned cell types based on the snRNAseq profiles using unbiased classifiers followed by manual curation (see Methods). This analysis captured the diversity of cell types expected based on the histopathology of cHL, including myeloid, lymphoid, and stromal cell types (fig. 1b, c). Given their importance and low abundance, HRS cells were identified using multiple orthogonal strategies. First, we identified putative HRS cells using the well-established marker genes *TNFRSF8* (CD30), *CD274* (PD-L1), *PAX5*, and *IRF4*. Next, we performed copy number variant inference to determine if this putative HRS cell population harbored shared copy number variation seen in clonal tumor cells but not normal cells. Strikingly, the putative HRS cell population showed strong evidence of copy number variation, whereas the other cell types did not, indicating that we had appropriately identified HRS cells (fig. 1d, e). EBV-positive HRS cells (as determined by in situ hybridization using EBV transcript-specific probes) were associated with higher levels of EBV transcript detection by RNAseq, supporting the high quality of the dataset (fig 1d). Notably, mapping RNA reads from EBV-negative HRS cells to pathogen reference databases did not reveal a signal suggestive of other pathogens playing a role in cHL (see Methods). Intriguingly, EBV-negative HRS cells tended to express expression programs typically seen in neuroglial cells (fig. 1f, extended data fig. 3a, b). The mechanism explaining this distinct transcriptional program requires further investigation.

Using the spatial profiling data, we next performed probabilistic cell-type mapping using the RCTD method^28^ to define the topology of HRS cells in tissues, validating the results by spatial plotting of marker genes, copy number variant inference, and EBV read mapping (extended data fig. 3c-f). Strikingly, in most tumors examined, the HRS cells showed a clustered spatial distribution (fig. 1g, extended data fig. 3g, supplementary table 4). This co-localization could reflect evidence of positive selection for paracrine HRS-HRS cell interactions.

### Defining the immune microenvironment in cHL

We next focused on the immune and stromal cells, which comprise the vast majority of cells within cHL tumors. We identified the presence of 23 distinct immune cell types (fig. 2a, extended data figs. 4a, b, 5a, b, e, f, 6a, b) and used compositional analysis^30^ to compare their abundance in cHL to that observed in reactive lymph nodes. We observed a striking increase in the abundance of activated and differentiated T cell subsets, including regulatory T cells, activated/proliferating CD4+ T cells, exhausted CD8+ T cells, and a concomitant decrease in less differentiated naive T cells. Moreover, monocytes/macrophages were significantly expanded, particularly in EBV+ cHL (fig. 2b, extended data fig. 4c, 5c, d, h, I; 6c, d).

**Fig. 2:**
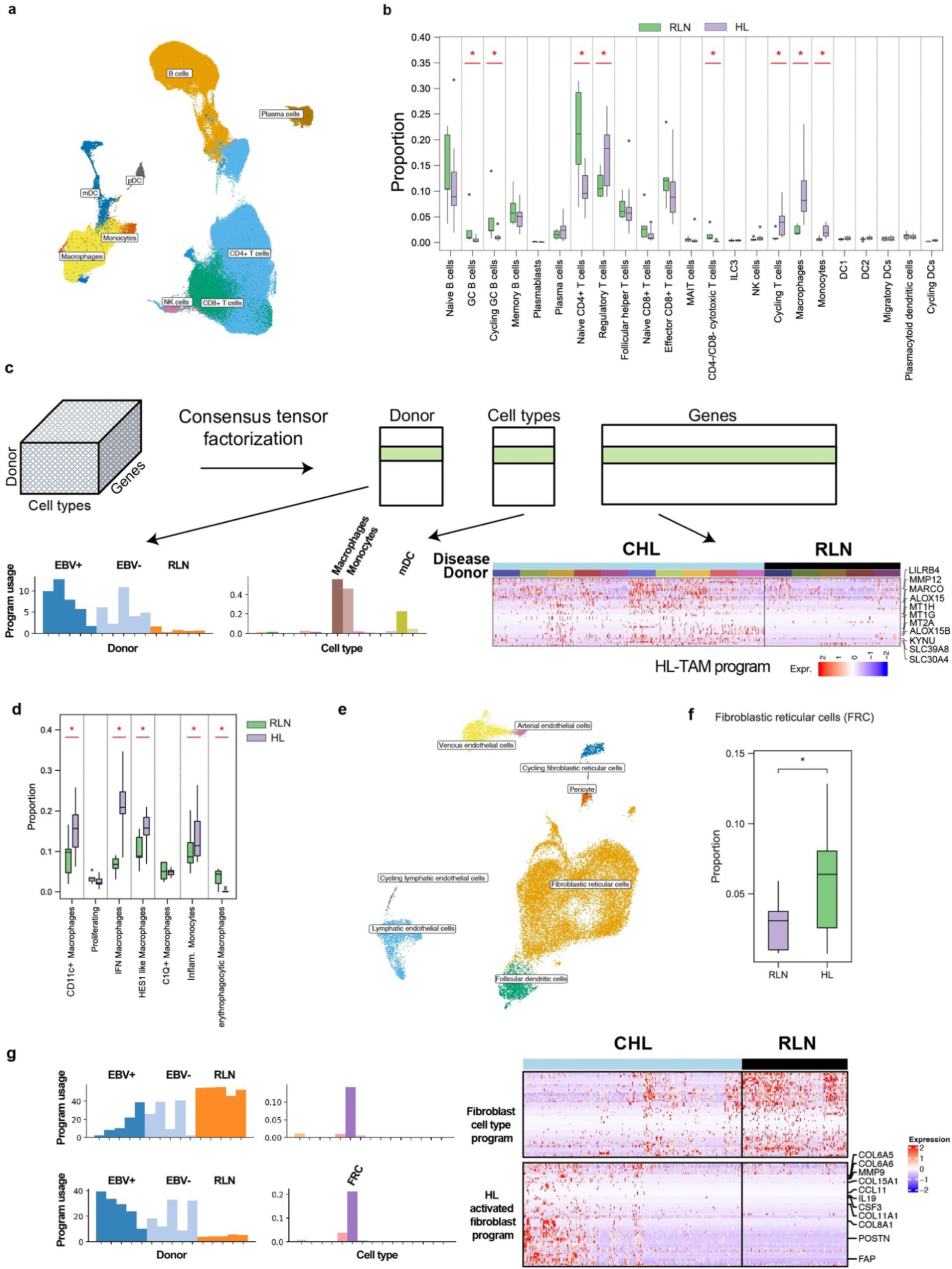
Compositional and transcriptional remodeling of the immune microenvironment in Hodgkin lymphoma. **a**, UMAP embedding of immune cells colored by cell type annotation. **b**, compositional analysis of immune cells in Hodgkin lymphoma (HL) versus reactive lymph node (RLN) samples. **c**, unsupervised consensus tensor factorization reveals a shared HL-associated myeloid cell program. **d**, compositional analysis of myeloid subsets across HL and RLN samples. **e**, UMAP embedding of stromal cells colored by cell type annotation. **f**, compositional analysis of stromal cells in HL versus RLN samples. **g**, consensus tensor factorization reveals a distinct activated pro-fibrotic, myofibroblast-like fibroblastic reticular cell program.

Beyond differences in cell type abundance, we observed substantial transcriptional changes within the immune cells. For example, cHL-associated germinal center B cells showed increased expression of granzyme B^31^ (GZMB; extended data fig. 4d), and cHL-associated regulatory T cells showed increased expression of IFNg-stimulated genes (IL12RB2, CXCL10) and NRN (neuritin), a secreted protein that enhances the expansion and suppressive function of Treg cells^32^ (extended data fig. 5g).

Unsupervised consensus tensor factorization analysis similarly revealed a distinct tumor-associated immunosuppressive myeloid cell program (fig. 2c) with coordinated upregulation of inhibitory receptors (MARCO, LILRB4), metal ion transporters (SLC39A8, SLC30A4), metal-binding proteins (MT1, MT2) and IFN and IL4/IL13 stimulated genes (ALOX15, KYNU). Further annotation of the monocyte/macrophage population revealed enrichment of inflammatory monocytes^33^ and the recently described^34^ IL4I1+/IFN-primed and HES1+ macrophages (fig. 2d, extended data fig. 6e,f). The enrichment of IL4/IL13 stimulated genes in myeloid cells is particularly notable as it suggests ongoing IL4/IL13 signaling in the tumor microenvironment.

Eight distinct stromal cell types were also identified in cHL tumors (fig. 2e, extended data fig. 6a), including enrichment of fibroblastic reticular cells independent of EBV status (fig. 2f, extended data fig. 6b). These cells exhibited a pro-fibrotic program characterized by expression of FAP, POSTN, collagens, and the pro-fibrotic cytokines CCL11 and IL19 (fig. 2g, extended data fig. 6c) and likely mediate the extensive fibrosis that is a histopathologic hallmark of cHL tumors^10^.

Importantly, spatial mapping of the immune microenvironment (extended data fig. 8a, b) revealed evidence of a distinct microenvironmental niche surrounding HRS cells. Specifically, we observed the enrichment of CD4+ T cells, follicular dendritic cells, and monocyte/macrophages near HRS cells. In contrast, fibroblasts, plasma cells, and plasmacytoid dendritic cells were depleted from HRS cell neighborhoods (fig. 3a, b). Consistent with these observations, immune cells near HRS cells had a distinct gene expression program compared to those same cell types located further away, perhaps explained by HRS cell-derived cytokine gradients (fig. 3c, extended data fig. 8c-e, h). We confirmed the existence of an HRS cell niche by generating an independent spatial dataset consisting of 637,000 cells from 4 formalin-fixed paraffin-embedded cHL samples analyzed on the CosMx platform measuring 1,000 transcripts (fig. 3d, extended data fig. 2b-d, 9a, b). These results argue in favor of the existence of a highly structured HRS cell niche as opposed to a random assortment of immune and stromal cell types (fig. 3e).

**Fig. 3:**
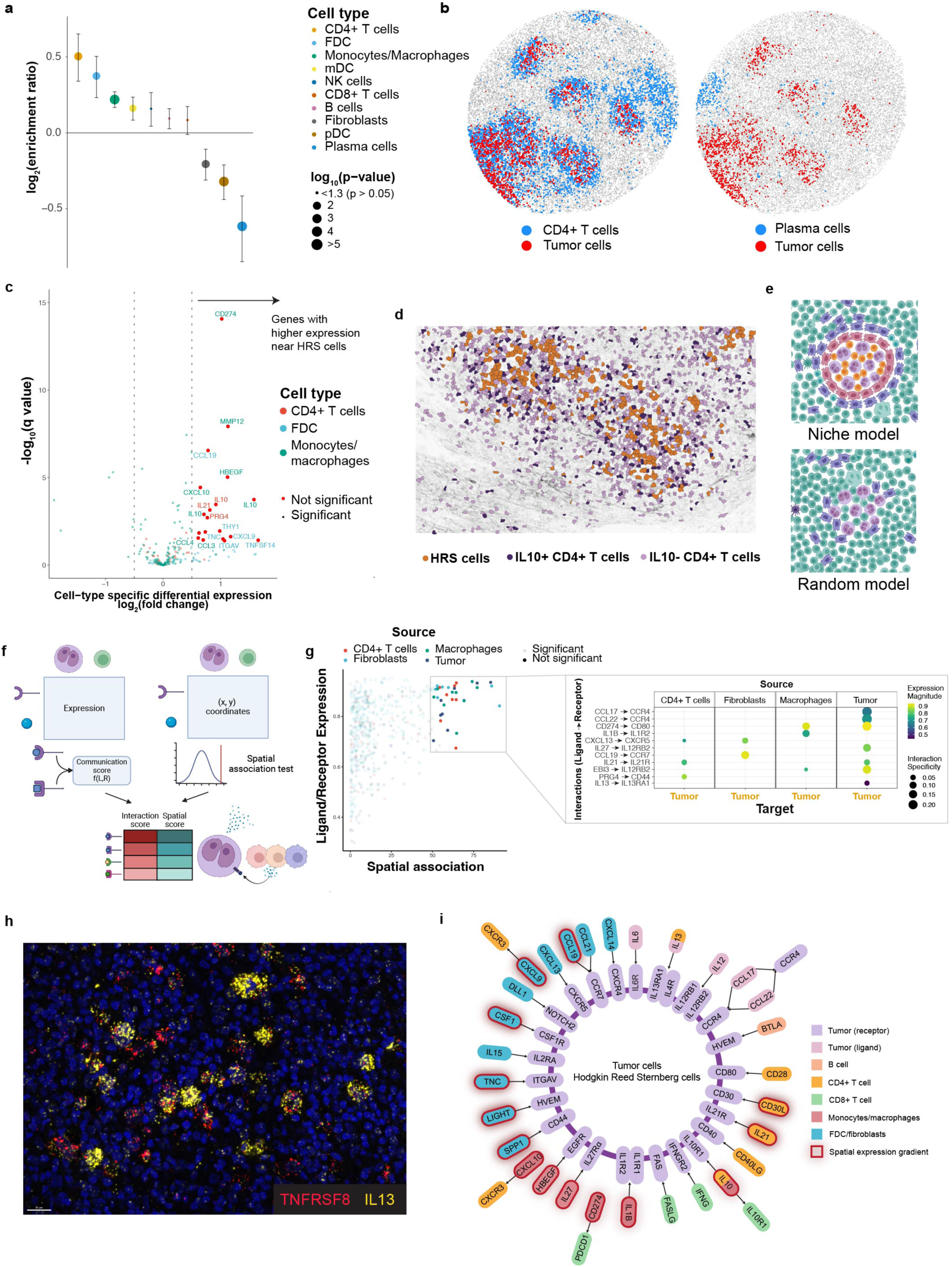
Spatial organization of the immune microenvironment around Hodgkin Reed Sternberg cells. **a**, Enrichment ratios of immune cell types (colored by cell type) aggregated across the entire multi-sample, multi-replicate slideseqV2 dataset. The size of the points indicates the p-value; mean ± s.e.m. is depicted. **b**, representative array showing spatial enrichment of CD4+ T cells and depletion of plasma cells around clustered HRS cells. **c**, volcano plot showing cell-type specific differentially expressed genes in relation to HRS cell proximal/dense regions. A positive log_2_(fold change) (right of volcano plot) indicates genes that show higher expression in HRS cell proximal/dense regions. Results are aggregated across the entire multi-sample, multi-replicate slide-seqV2 dataset. **d**, CosMx analysis showing enrichment of IL10+ CD4+ T cells within HRS cell dense/proximal regions. **e**, schematic representation of the structured model where immune cells show a non-random spatial distribution around clustered tumor cells in contrast to the null hypothesis random spatial model where immune cells are randomly distributed. **f**, schematic overview of the approach for systematic discovery and prioritization of spatially-informed tumor-immune ligand-receptor interactions within the HRS cell niche. **g**, scatterplot of tumor-targeted ligand-receptor interactions based on spatial proximity (spatial association metric, see methods) and normalized ligand/receptor expression (scaLRscore metric). Interactions that show high ligand/receptor expression and spatial proximity are called out and shown on the right as a bubble plot. **h**, RNA fluorescence in-situ hybridization analysis of IL13 expression (yellow) in HRS cells and the microenvironment. The red TNFRSF8 (encoding CD30) probe highlights HRS cells. Scale bar; 20 µm. The data are representative of independent experiments on three different patient samples. **i**, schematic diagram of systematically prioritized paracrine tumor-microenvironment interactions in Hodgkin lymphoma, including putative growth factors and immunosuppressive interactions

### Identification of ligand-receptor interactions in the HRS cell niche

We next sought to identify secreted or cell surface ligands expressed by microenvironmental cells in proximity to HRS cells. We further prioritized such survival factors by requiring their receptors to be expressed in HRS cells. This spatially-aware ligand-receptor analysis revealed multiple candidate immunoregulatory and growth factor interactions validated in the independent dataset and via in situ hybridization (fig. 3f-I, extended data figs. 8f, g, and 9c-n). This analysis recovered known interactions involving CD274 (PD-L1), CCL17, and CCL22 ligands, highlighting the effectiveness of the approach.

### Experimental validation of candidate juxtacrine and paracrine growth factor interactions

Having systematically identified candidate spatially associated ligand-receptor interactions that might support HRS cell growth or survival, we sought to test the importance of these associations experimentally. We first performed gain-of-function experiments, asking whether any of the 36 tested recombinant candidate ligands increased the abundance of the HRS cell lines L1236 and UHO1 when grown in either low serum or low-density conditions. 20/36 (55%) yielded statistically significant increases in cell numbers, with the greatest effect seen with CD40 ligand (CD40LG) and IL13 (fig. 4a and extended figs. 10a, b).

**Fig. 4:**
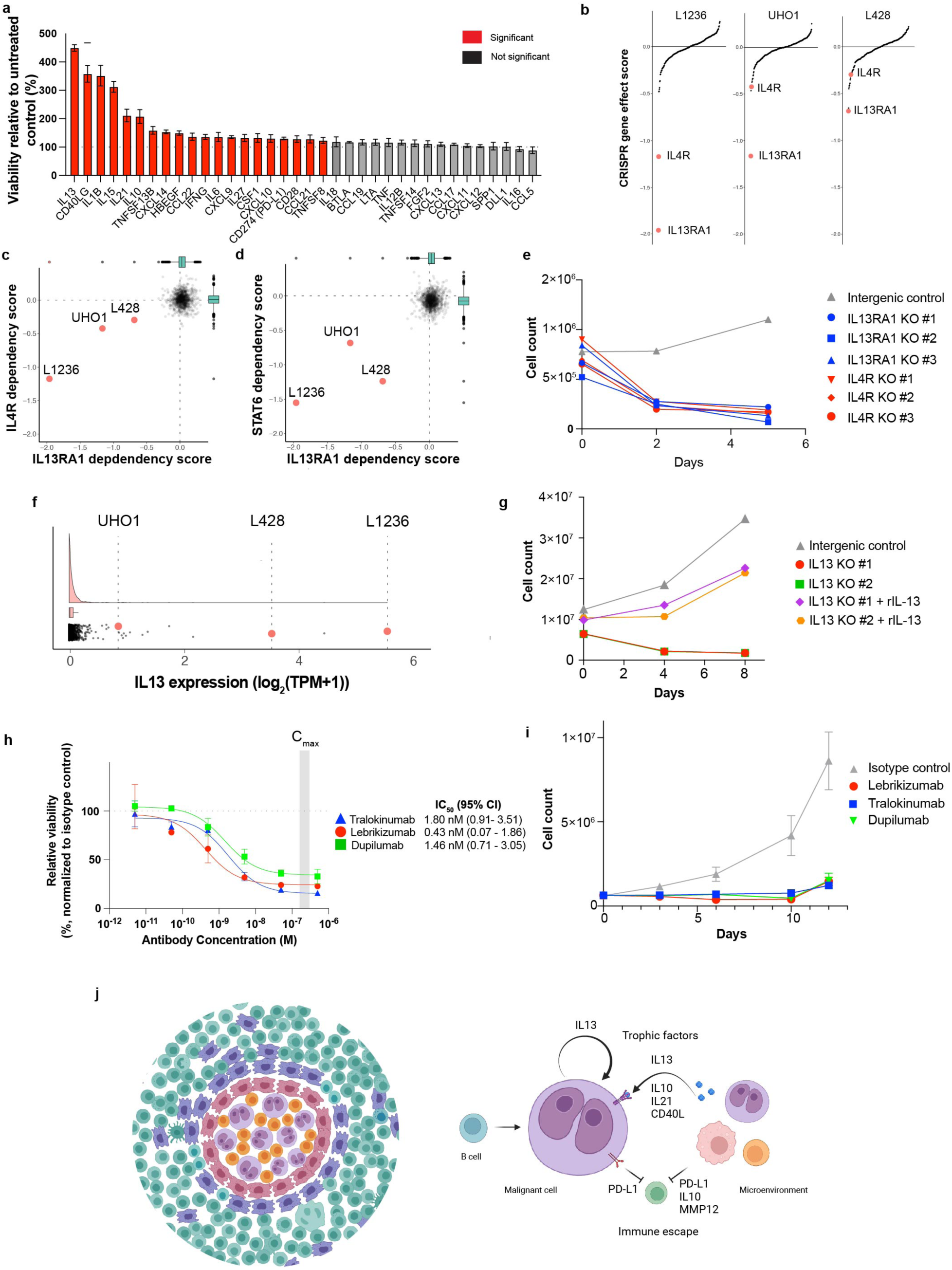
Experimental validation of paracrine growth factors for Hodgkin-Reed Sternberg cells. **a**, Barplot showing the effect of recombinant factors on the relative viability (normalized to untreated control) of L1236 cells at three days under low serum conditions; mean ± s.d. is depicted (n = 3); *P* < 0.05 is considered significant; two-way analysis of variance (ANOVA) followed by Dunnett’s multiple comparison test. **b**, waterfall plots (ordered by increasing CRISPR gene effect score) showing genetic dependencies of 3 cHL cell lines (L1236, UHO1, L428). **c, d**. scatterplot of IL4R and IL13RA1 (c) and IL4R and STAT6 (d) gene effect scores across the DepMap dataset. **e**, effect of CRISPR/Cas9-mediated knock out of IL13RA1 or IL4R on L1236 cell viability (viable cell counts). **f**, rainplot of IL13 expression showing that the cHL cell lines (L1236, L428, and UHO1) show outlier expression of IL13. **g**, effect of CRISPR/Cas9-mediated knock out of IL13 on L1236 cell viability (viable cell counts), with or without recombinant IL13 supplementation (rIL13). **h**, dose-response analysis of tralokinumab, lebrikizumab, and dupilumab on L1236 cells. IC_50_ values (95% confidence interval) are indicated on the right (n = 2); mean ± s.d. is depicted. The relative viability values represent the mean of two independent experimental replicates. **i**, time course analysis of the effect of tralokinumab, lebrikizumab, dupilumab (0.5 µM each), and isotype control antibody on L1236 cell viability (n = 3); mean ± s.d. is depicted. The C_max_ range of these antibodies reported in clinical trials is indicated in grey. **j**, schematic representation of the proposed model of spatial organization and interactions that mediate tumor cell growth within the intact cHL tumor microenvironment. The data in e, g, and i are representative of three independent experiments.

To test whether IL13 and CD40LG signaling was essential for the survival of HRS cells, we first turned to the Cancer Dependency Map (DepMap)^35^, a compendium of 1,086 human cancer cell lines subjected to genome-wide CRISPR/Cas9 knock-outs. Whereas CD40 was not a dependency in any of the DepMap models, knock-out of either component of the obligate heterodimeric receptor for IL13 (IL4R and IL13RA1) resulted in a profound loss of HRS cell viability. Furthermore, STAT6, the downstream transcriptional mediator of IL13 signaling, scored as an HRS cell dependency. Specifically, all three HRS cell lines in the DepMap required IL4R, IL13RA1, and STAT6 expression for survival, whereas none of the other cell lines in the database exhibited such dependency (fig. 4b, c, d). This finding is extremely unlikely to be seen by chance alone (IL13RA1, *P* = 4.8×10^-9^; IL4R, *P* = 5.1×10^-6^; STAT6, *P* = 4.7×10^-9^; Fisher’s Exact Test of Independence). We further validated this result using independent, individual gRNAs targeting IL4R and IL13RA1 and showed they result in loss of HRS cell viability (fig. 4e). Analysis of these cells showed induction of cleaved caspase 3, consistent with cell death via apoptosis (extended data fig. 10c). We hypothesize that these HRS cell lines underwent selection for their ability to express IL13, a cytokine that is not typically expressed at high levels by normal B cells^35^. Consistent with this hypothesis, the HRS cell lines showed outlier expression of IL13 (fig. 4f), and knock-out of IL13 similarly resulted in loss of cell viability that was rescuable with recombinant IL13 (fig. 4g, extended data fig. 10d).

We next asked whether the genetic dependency of IL13 signaling in HRS cells could be phenocopied pharmacologically. Specifically, we tested three monoclonal antibodies that block IL13 signaling through distinct mechanisms: dupilumab, which targets the alpha subunit of IL4 receptor (IL4R); tralokinumab, which blocks the binding of IL13 to IL13RA1 and lebrikizumab, which impairs the heterodimerization of IL4R and IL13RA1. An isotype control antibody served as a negative control. All three antibodies showed dose-responsive and time-dependent loss of viability of HRS cells and modest induction of apoptotic cell death, whereas the isotype control showed no effect (fig. 4h, i, extended data fig. 10e, f). These results are particularly notable because dupilumab, tralokinumab, and lebrikizumab are FDA-approved for use in patients with atopic dermatitis^36–39^. Importantly, we observed maximal growth inhibitory activity in HRS cells at concentrations that are readily achievable in patients treated at the clinically approved dose (fig. 4h, extended data fig. 10e).

## Discussion

Anatomical pathologists have long recognized that tumors are complex admixtures of malignant cells and normal cell types. While hypothesis-driven research on the tumor microenvironment has yielded important insights, systematic genomic approaches have largely been restricted to the analysis of tumor cells. Researchers have typically gone to great lengths to purify cancer cells away from the microenvironment to enrich the signal coming from the malignant component of the tumor. More recently, single-cell transcriptomics has allowed for the analysis of all cells within a tumor, but the spatial information is lost with such approaches, thereby precluding the analysis of interactions between tumor cells and the normal cells immediately surrounding them.

Hodgkin lymphoma is one of the most enigmatic human tumors, with classic Hodgkin lymphoma (cHL) being composed of only rare tumor cells surrounded by an extensive array of infiltrating immune and stromal cells. While it is possible that these microenvironmental cells are unimportant for tumor cell survival, it has been well-documented that removing the malignant HRS cells from the microenvironment results in their failure to survive in vitro^13–15^. We, therefore, hypothesized that the microenvironment is a source of such survival and immune evasive factors.

To address this, we generated the most comprehensive single nucleus and spatial profiling dataset to date, composed of more than 2.2 million transcriptional profiles. Importantly, we believe this reference dataset includes all relevant cell types within cHL tumors, whereas standard single-cell profiling methods^25,26,40,41^ result in the loss of key cell types due to limitations of cell isolation methods. For example, macrophages do not survive isolation, yet they can comprise more than 50% of cells within cHL^27^. The quality of this reference dataset is supported by multiple lines of evidence, including the demonstration of clonality of rare HRS cells, the matching of cell types recognized by immunohistochemical analyses, and the confirmation of the recently described^32,41^ expansion of LAG3+ regulatory T cells and IL1B+ inflammatory monocytes that are thought to contribute to the ability of cHL to avoid immune surveillance.

Our study complements the growing body of literature supporting the presence of a distinct pro-tumorigenic cellular niche around HRS cells in cHL. Consistent with previous work, we observed the enrichment of CD4+ T cells, monocytes/macrophages, and myeloid dendritic cells and the depletion of plasmacytoid dendritic cells near HRS cells^23,42,43^. We additionally show that plasma cells are excluded from the immediate proximity of HRS cells, which may represent a potential mechanism of evading the humoral immune response. Previous hypothesis-driven studies^23,41–43^ have shown that specific immune checkpoint molecules are expressed in the cHL niche, including the myeloid markers (PD-L1, TIM3, IDO1) and CD4+ T cells (PD1, CTLA4, LAG3). Our genome-wide analysis recovers the recently reported^44^ interaction between CXCR5+ HRS cells and CXCL13-expressing macrophages and extends the observation to CXCL13-expressing CD4+ T cells, follicular dendritic cells and fibroblasts in proximity to HRS cells.

We experimentally validated one of the strongest findings of the cHL microenvironment, namely the nomination of IL13 as a potential survival factor for HRS cells. Gain- and loss-of-function studies supported the IL13 hypothesis. Analysis of the Cancer Dependency Map^34^ showed that IL4R and IL13RA1 were required for the survival of only three cell lines in the DepMap, and these are the only three cHL cell lines in the dataset. The only other gene with this pattern of selective dependency is STAT6, known to be regulated only by IL4 and IL13 signaling^45–47^. Of note, IL13 itself does not score in the DepMap dataset because of the pooled nature of those experiments; knock-out of IL13 would be rescued by nearby IL13-expressing cells. However, non-pooled experiments showed that IL13 is indeed required for the survival of cHL cell lines.

We note that CD4+ T cells in the cHL microenvironment express IL13, consistent with IL13 serving as a key T cell cytokine. However, consistent with previous reports^48,49^, we also observed robust IL13 expression in HRS cells themselves – a surprising phenomenon given that B cells typically do not express high levels of IL13^35^. We postulate that surrounding T cells may establish IL13 signaling dependency in the early stages of cHL evolution, with subsequent selection for paracrine secretion of IL13 by HRS cells either by epigenetic or somatic genome alterations. The recent description of somatic activating mutations in IL13RA1 is consistent with this hypothesis, further supporting that HRS cells invoke multiple mechanisms to ensure IL13 signaling^50^.

To explore the clinical translational potential of the role of IL13 signaling in cHL, we tested three IL13 receptor-blocking antibodies for their ability to recapitulate the effect of IL13, IL4R, or IL13RA1 genetic knock-out in cHL cell lines. In all three cases, antibody treatment at clinically achievable doses resulted in growth inhibition and apoptotic cell death. This finding is particularly noteworthy because these antibodies are already FDA-approved for treating atopic dermatitis^36–39^. We also note the recent report of using one of these antibodies, Dupilumab, to enhance responsiveness to PD1/PD-L1 checkpoint inhibitors in patients with lung cancer^51^. Taken together, these results suggest that IL13-blocking antibodies should be evaluated for their activity in patients with cHL. Our results also indicate that the cHL tumor microenvironment is a rich source of opportunities for therapeutic intervention, and the availability of a cHL spatial reference dataset should catalyze the evaluation of such targets. More broadly, our study describes a general approach for discovering cellular niches and microenvironment-derived therapeutic targets.

## Methods

### Tissue collection

After appropriate institutional review board approval and informed consent at the Brigham and Women’s Hospital, Massachusetts General Hospital, and the Broad Institute, excess excisional biopsy tissue specimens were collected from patients with newly diagnosed classic Hodgkin lymphoma. Additionally, reactive lymphoid tissue control samples were also collected. Demographic information for all study participants is provided in Supplementary Table 1. The specimens were embedded in OCT compound, snap-frozen, and stored at -80°C. As a quality control step, tissue architecture was assessed by hematoxylin and eosin staining, and RNA integrity was determined using the Tapestation RNA ScreenTape system (RINe > 7.5).

### Single nucleus isolation and RNA sequencing

Fresh frozen tissues (embedded in OCT) were cryo-sectioned at 20-micron thickness on a Leica CM1950 Cryostat (Leica, US) at -20°C. Pre-cooled 3 mm circular (3332P/25, Integra) biopsy punches were used to isolate regions of interest from tissue sections. The punched tissue sections were placed in wells of a 6-well plate and 1mL of extraction buffer (82 mM Na_2_SO_4_, 30 mM K_2_SO_4_, 10 mM Glucose, 10mM HEPES, 5 mM MgCl_2_, 1% Kollidon VA64, 1% Triton X100, 0.1% BSA, 667 units/mL RNase-inhibitor (Biosearch technologies, 30281-1)) was added to each well. The sample was then triturated slowly using a P1000 24 times over 2 minutes, repeated five times. After dissociation, the tissue was transferred to a falcon tube, and the volume was diluted to 30 mL with a wash buffer (82 mM Na2SO4, 30 mM K2SO4, 10 mM Glucose, 10mM HEPES, 5 mM MgCl2, 0.1% BSA, 67 units/mL RNase-inhibitor. Diluted samples were split into two 50 mL falcon tubes and centrifuged at 600 rcf for 10 min. Most of the supernatant was removed, and the samples were repooled. Pooled samples were passed through a 40 μm strainer and collected in a pre-cooled 1.5 mL Eppendorf tube. In a pre-cooled centrifuge, samples were further concentrated for 10 min at 200 rcf. All but 50 μL of the supernatant was discarded, and the pellet resuspended. This solution was stained with DAPI (Thermo Fisher Scientific, 62248), and the nuclei were quantified using a C-Chip Fuchs-Rosenthal disposable hemocytometer (INCYTO, DHC-F01-5). The sample was then appropriately diluted to recover 10,000 cells according to the 10x Genomics 3’ gene expression (v3.1 chemistry) user guide (CG000204, Rev D). Partitioning oil and gel beads were added to the Chromium Next GEM Chip G with the single nuclei suspensions. Following successful GEM generation, reverse transcription was performed, and cDNA was recovered and cleaned up using Dynabeads MyOne SILANE according to the 10x Genomics protocol. The recovered cDNA was then amplified for 12 cycles, and the products were quantified using Agilent Bioanalyzer High Sensitivity DNA Chip. Once cDNA quality was confirmed to be within expectations, the libraries were prepared according to the Chromium Next GEM Single Cell 3’ Reagent Kits v3.1 steps. Following the 10x Genomics library preparation, the samples were qualified using the High Sensitivity DNA Chip and quantified using the Qubit 3.0 fluorometer. The libraries were normalized to 2 nM and pooled. Libraries were sequenced on an Illumina Nextseq 550/500 or a Novaseq 6000 S2 flow cell and demultiplexed for downstream analysis.

### Spatial transcriptomics

#### Slide-seqV2

Alternate 10 μm thick tissue sections were thawed onto 3mm Slide-seq arrays. The arrays were transferred immediately to a hybridization solution (6X sodium chloride sodium citrate (SSC) with 2 U/μL Lucigen NxGen RNAse inhibitor) for 30 minutes at room temperature. The intervening sections were used for hematoxylin and eosin, as well as immunohistochemical staining. Libraries were generated using the previously published Slide-seqV2 protocol ^52,53^. The libraries were prepared using the standard Illumina protocol and sequenced on either Nextseq 550/500 or Novaseq 6000 S2/S4 flowcells at a depth of 100 million reads per array. Samples were pooled to a concentration of 4 nM and were sequenced with the previously described read structure^53^. The final dataset includes at least two biological replicates (multiple Slide-seq arrays) per patient sample.

#### Visium Spatial Gene Expression for FFPE V1

Two formalin fixed paraffin embedded samples of newly diagnosed classic Hodgkin lymphoma (from two unique patients) were selected based on the DV200 metric (> 50% of RNA molecules greater than 200 base pairs in length) determined using the Tapestation RNA ScreenTape system. Two 5-micron sections from each tumor were placed on the four 6.5 x 6.5 mm capture areas of the Visium spatial gene expression slide. Deparaffinization, staining (Hematoxylin and Eosin), imaging, de-crosslinking, overnight probe hybridization, ligating, barcoding, and library construction was performed per manufacturer recommendations (protocol CG000404 Rev B). Quality control analysis of libraries was performed using the Tapestation system, and verified libraries were sequenced on a Nextseq 500 high output flow cell at a depth of >10,000 reads per spot for sample A and >45,000 reads per spot for sample B, which approaches saturation (>80%) for both samples.

#### CosMx Spatial Molecular Imager

Formalin-fixed, paraffin-embedded (FFPE) tissue sections were prepared for CosMx SMI profiling as previously described^54^. NanoString® ISH probes were prepared by incubation at 95°C for 2 min and placed on ice. The ISH probe mix (1nM 980 plex ISH probe, 10nM Attenuation probes, 1X Buffer R, 0.1 U/μL SUPERase•In™ [Thermofisher] in DEPC H2O) was pipetted into the hybridization chamber. The hybridization chamber was sealed to prevent evaporation, and hybridization was performed at 37°C overnight. Tissue sections were rinsed of excess probes in 2X SSCT for 1 min and washed twice in 50% formamide (VWR) in 2X SSC at 37°C for 25 min, then twice with 2X SSC for 2 min at room temperature and blocked with 100 mM NHS-acetate in the dark for 15 min. A custom-made flow cell was affixed to the slide in preparation for loading onto the CosMx SMI instrument.

RNA target readout on the CosMx SMI instrument was performed as described previously^54^. After RNA readout, the tissue samples were incubated with a 4-fluorophore-conjugated antibody cocktail against CD298/B2M (488 nm), PanCK (532 nm), CD45 (594 nm), and CD3 (647 nm) proteins and DAPI stain in the CosMx SMI instrument for 2 h. After unbound antibodies and DAPI stain were washed with Reporter Wash Buffer, Imaging Buffer was added to the flow cell, and eight Z-stack images for the 5 channels (4 antibodies and DAPI) were captured.

### Single-molecule RNA fluorescence in-situ hybridization

The RNAscope HiPlex V2 assay (ACDbio) was performed as previously described. FFPE sections were rehydrated on ice for 3 hours before sectioning. Tissue was cut to 5 μm sections and placed on charged slides. Slides were air-dried at room temperature overnight. The following day, slides were incubated at 60°C for 2 hours. Tissue sections were deparaffinized and rehydrated in xylene, followed by ethanol and water per Advanced Cell Diagnostics (ACD) recommendations.

### Histopathology and immunohistochemistry

#### Hematoxylin and Eosin (H&E) staining

Frozen tissue sections on glass slides were dipped in xylene, hydrated with a graded ethanol wash series, and stained with hematoxylin. A weakly alkaline ammonium hydroxide solution was used as a nuclear “bluing” reagent. The sections were stained with eosin and dehydrated through a graded ethanol series, xylene, dehydrated and coverslipped. Brightfield images were taken using the Leica Aperio VERSA Brightfield, Fluorescence & FISH Digital Pathology Scanner under a 10x objective.

#### Immunohistochemistry

Immunohistochemical staining of frozen and formalin-fixed paraffin-embedded tissue sections was performed according to clinical procedures in the Department of Pathology at the Brigham and Women’s Hospital, as previously described^55^.

### Functional validation assays

#### Cell culture

Mammalian cell lines L428, L1236, and UHO1 were obtained from the Leibniz Institute DSMZ and cultured in media and conditions recommended by the distributor. After initial thawing and one month of passaging, cells were sent to LabCorp for STR Profiling and Mycoplasma testing. Cells were regularly tested for Mycoplasma contamination using the Mycoscope PCR Mycoplasma Detection Kit (BIO-CAT MY01050-GL).

#### Pharmacologic assays

We used relative viability measurements and total cell counts to determine the dose-dependent response and time-to-effect of three IL13-blocking antibodies. For these experiments, the following three antibodies were used: Tralokinumab (MedChemExpress HY-P99053), Dupilumab (Selleck Chemicals A2038-1mg), and Lebrikizumab (MedChemExpress HY-P99025). Human IgG4 Lambda (Millipore Sigma I4765-1MG) and Kappa (MedChemExpress HY-P99003) were used as isotype controls.

Each therapeutic antibody or isotype control was serially diluted (0.5 µM to 50 pM) in 96-well plates. Then, 20,000 cells were added per well and cultured for nine days. On day nine, relative viability was assessed using the CellTiter-Glo (CtG) Luminescent Cell Viability Assay (Promega G7572) per the manufacturer’s instructions. Luminescence was quantified using an EnVision Multilabel Plate Reader, and each antibody-treated condition was normalized to its isotype control. For the time course experiments, all antibody and isotype control treatments were 0.5 µM. At regular intervals, viability measurements were made using CtG and viable cell counts (Vi-cell Blue). The culture media was changed every three days, and the antibody was replenished.

#### Cytokine supplementation assays

To investigate the effect of putative growth factors identified computationally, L1236 and UHO1 cells were supplemented with recombinant cytokines (concentrations and sources in Supplementary Table 6). Cytokines were reconstituted and aliquoted according to the manufacturer’s recommendations. The assays were performed in 96 well plates using 20,000 cells per well in triplicate. Cytokine was replaced during media changes every three days, and viability was measured using CtG.

#### CRISPR/Cas9 knockout

Kanamycin-resistant plasmids containing the top three CRISPR/Cas9 guides for IL13, IL13Ra1, and IL4R (Supplementary Table 7) were obtained from the Genomic Perturbation Platform. After overnight culture of E. coli in LB medium (50 µg/mL Kanamycin), plasmid DNA was extracted with the QIAGEN Maxi Plasmid Purification kit (Cat. 12162). Lentiviral particles were produced in HEK293T cells by co-transfection of guide plasmids with psPAX2 (Addgene #12260) and pMD2.G (Addgene #12259), then concentrated using Lenti-X Concentrator (Takara PT4421 2). L1236 cells engineered to express Cas9 (with Blasticidin and Puromycin resistance) were maintained in standard conditions plus 4 µg/mL Blasticidin S HCl (Invitrogen A1113903) for two weeks to ensure Cas9 activity. For transduction, 10 million cells per condition (plus a non-transduced control) were seeded in 6-well plates. The following day, Polybrene (Millipore-Sigma TR-1003-G) was added to 8 µg/mL, and 300 µL of concentrated lentivirus was added to each well (or DMEM to non-transduced controls). Cells were incubated overnight, then media was replaced, and 2 µg/mL Puromycin Dihydrochloride (Invitrogen A1113803) was added to select transduced cells. Knockouts targeting IL13 were supplemented with 0.1 µg/mL of recombinant IL13 (PeproTech 200-13). Cells were counted regularly for 15 days on the Vi-cell Blue. Once the population neared the instrument’s lower detection limit, genomic DNA was extracted (QIAamp DNA Micro kit, QIAGEN 56304). Editing efficiency (>75% edited sequences) was confirmed by CRISPR sequencing (MGH CCIB DNA core; CRISPRESSOv2 analysis; Supplementary Table 8), as no high-quality antibodies were available to verify protein knockout.

#### Apoptosis assays

Caspase 3/7 activity was measured 24 hours after antibody treatment or transduction/selection using the CellEvent green fluorescence assay per manufacturer recommendations. Hoechst 33342 was used for nuclear counterstaining. The assay was performed with at least three technical replicates. Imaging was performed using the EVOS M7000 imaging system using the DAPI and GFP channels. The mean nuclear intensity of CellEvent green fluorescence was calculated per replicate using CellProfiler and normalized to the baseline level of fluorescence observed with the intergenic control. Staurosporine (1μM) was used as positive control.

#### Dependency Map data analysis

Gene effect scores derived from CRISPR knockout screens published by Broad’s Achilles^35^ and Sanger’s SCORE^56^ projects were downloaded from the internal DepMap portal (public 22Q2). Gene effect scores were inferred by Chronos^57^, with negative scores implying a fitness defect following gene knockout. As previously described, the scores are normalized such that non-essential and common essential genes have a median score of 0 and -1, respectively. A cell line was considered dependent if the probability of dependency was > 0.5. The probability of dependency is calculated for each gene effect score in a cell line as the probability that the score arises from a distribution of essential gene scores rather than non-essential gene scores^58^. To test whether any of our predicted ligand-receptor interactions were dependencies in Hodgkin lymphoma cell lines, we focused our analysis on our prioritized list of ligands and receptors and excluded common essential genes. We performed a Fisher’s exact test of independence to determine if there was any association between dependency on these genes (IL13RA1, IL4R, STAT6) and cancer type (Hodgkin lymphoma versus other cell lines).

### Single nucleus RNA sequencing data analysis

#### Count matrix generation

Raw BCL files were demultiplexed and converted to FASTQ, which were then aligned to the GRCh38 human genome reference (cellranger reference 3.0.0, Ensembl v93 gene annotation, introns included) using the Cellranger count and mkfastq commands that are implemented in the Cumulus ^59^ Cellranger workflow (v2.3.0) on Terra.

#### Ambient RNA correction and excluding empty droplets

We used the ‘remove-background’ function of CellBender v0.2.0^60^ as implemented in the Cumulus CellBender workflow (v2.1.1) on Terra. The parameter ‘expected-cells’ was obtained from the Cell Ranger metric ‘Estimated Number of Cells’.

#### Doublet prediction

The gene-barcode expression count matrices were processed individually in R using Seurat v4. Initial filters were applied to exclude genes expressed in fewer than 3 cells and cells with 200 or fewer genes. ScDblFnder^61^ was then used to estimate doublets for each sample individually, with the expected number of doublets set as 1% per 1000 cells captured. The uncorrected expression count matrix (before ambient RNA correction using CellBender) was used as the input for doublet prediction.

#### Quality control, filtering, and dimensionality reduction

All the individual sample datasets were merged into one combined dataset. Filters were applied to exclude all estimated doublets, cells with 200 or fewer genes, 400 or fewer transcripts, and % mitochondrial reads > 5%. The combined, filtered count matrix was log-transformed and normalized using the NormalizeData function and the LogNormalize method. The top 2000 highly variable genes were selected for downstream analysis using the ‘vst’ method in the FindVariableFeatures function. The expression matrix was then scaled and centered using the ScaleData function. We then performed dimensionality reduction using the principal component analysis (PCA) and further performed uniform manifold projection and approximation (UMAP) using the first 50 principal components. The UMAP embeddings were used only for visualization purposes.

#### Data Integration

To facilitate cell type annotation, we used Harmony v0.1^62^ to construct a shared embedding in which cells group by shared cell types across multiple samples while minimizing technical sources of variation. We performed integration using multiple covariates that reflect multiple partially related sources of biological and technical variation in the data (sequencing batch, library preparation batch, sequencing technology, donor, and GEM lane); see supplementary table 1 for complete details of batch variables. Convergence was achieved in 10 iterations, and the first 50 Harmony-corrected PCs were used to compute UMAP embeddings and a shared nearest-neighbor graph for downstream analysis and visualization.

#### Cell type annotation

Preliminary cell type labels were obtained by applying the SingleR algorithm using multiple immune cell reference datasets (HumanPrimaryCellAtlasData, BlueprintEncodeData, DatabaseImmuneCellExpressionData, MonacoImmuneData). The CellTypist algorithm was also used to assign preliminary cell type identities using the Immune_All_Low and Immune_All_High references. Initial broad cell type labels were assigned manually using a combination of automated annotation results and known cell type markers/signatures of immune and stromal cell types. This approach led to the identification of tumor (HRS) cells, B/plasma cells, T/NK cells, myeloid cells, and stromal cells (fibroblasts and endothelial cells). Next, we split the Seurat object into these major cell types and re-ran the scaling, dimensionality reduction, and clustering procedure described above. The resulting clusters were annotated using the same approach, and the cell type labels were added to the main integrated object.

#### CNV inference

We use inferCNV v.1.7.1^63^ to identify evidence for somatic large-scale chromosomal copy number alterations in HRS cells by comparing gene expression intensity across positions of tumor genome in comparison to a set of reference ’normal’ cells in the single nuclei dataset. For this analysis, we subset the data to include samples from cHL patients only. We use the cell-type annotation described previously to select HRS cells as the malignant set and all other cell types as the ‘normal’ reference. Due to performance constraints, we downsampled the reference cells so that there were a maximum of 7,000 cells from each reference cell type. We use the “denoise,” “HMM,” and “cluster_by_groups” settings to enable the denoising procedure, CNV predictions via HMM, and separate clustering of cells from each patient, respectively. We set the cutoff value to 0.3 and the sd_amplifier to 2. All other settings were set to the default values. Gene expression levels were represented in a heatmap where genes were sorted by genomic position and ordered within each chromosome.

#### Cell-type compositional analysis

We use the single-cell compositional data analysis framework (scCODA)^30^ to compute the statistical significance of differences in cell-type composition between sample groups in the single-nuclei data. We specifically test for effects of the binary covariates *disease_status* (10 Hodgkin Lymphoma versus 5 Healthy samples) and *ebv_status* (5 EBV positive versus 5 EBV negative samples) in cell-type proportion obtained from *level 4* annotations. To run scCODA, we first compute the proportion of each of the cell types for each donor. We then use scCODA’s automatic reference selection to select a reference cell type with the least relative abundance dispersion over all samples while being present in at least 95% of the samples. The reference cell type selected for the single-nuclei data using this criterion was ILC3. We run scCODA’s parameter inference routine using the Hamiltonian Monte Carlo sampling method with the default chain length of 20,000. Following the guidelines in the scCODA package, we set an FDR of 0.2 for the *disease_status* analysis and an FDR of 0.4 for the *ebv_status* analysis. The main figures report the cell types for which the given covariate has a statistically credible effect.

#### Cell type-specific differential expression analysis

To identify genes differentially expressed between different sample groups within specific cell types in the single-nuclei data, we perform pseudo-bulk differential gene expression analysis using muscat^64^ and DESeq2. We first obtain pseudo-bulk expression profiles by summing the raw UMI counts for each gene, sample, and cell type using the aggregateData method in muscat. We filter out cell-type-sample pairs with less than 10 cells to ensure sufficient statistical power. All of muscat’s internal sample and gene filtering steps were also enabled. We then test differential expression using the DESeq2 method incorporated in muscat’s pbDS function. A gene was considered differentially expressed in a given cell type if the Benjamini-Hochberg adjusted local (within cell-type) p-value was < 0.05 and absolute fold change was > 2.

#### EBV reference mapping

For both 10x genomics single-nuclei and slide-seq spatial transcriptomics libraries, we used host genome (hg38) mapped BAM files as our starting point and extracted unmapped reads (flag 4 set) using samtools, which contain potential viral or otherwise non-host derived transcripts. To find EBV transcripts, we used NCBI reference sequence NC_007605.1. Several low-complexity regions of this genome would give rise to false positive alignments of low-quality reads. As such, we used USEARCH to mask these regions using the fastx_mask function. We mapped the unmapped reads using STAR aligner to this filtered genome. For 10x libraries sequenced with 90 cycles on read2, we used a stringent alignment score (tag ‘AS’) of 70 and number of mismatches (tag ‘nM’) of at most 4 to assign confident EBV calls. For slide-seq libraries sequenced with 40 cycles, we used an alignment score of 30 and number of mismatches of at most 2. Barcodes and UMIs associated with each confidently mapped read were then extracted from the original BAM file and deduplicated to find the number of EBV-mapped transcripts in each 10x droplet or slide-seq spatial spot.

#### Disease-associated immune cell signatures analysis

We use tensor factorization framework Consensus Zero Inflated Poisson Tensor Factorization (C-ZIPTF)^65^ to identify significant gene expression programs that can be associated directly with biological processes within specific cell types and donor contexts from the single-nuclei data in an unsupervised manner. We first create a pseudo-bulk expression tensor by summing the raw UMI counts for each gene, donor, and cell type. The pseudo-bulk data is represented as a 3-way tensor χ with dimensions *N*_*D*_ × *N*_*C*_ × *N*_*G*_, where *N*_*D*_, *N*_*C*_, and *N*_*D*_ represent the number of donors, cell types, and genes, respectively. χ _*i,j,k*_ denotes the aggregate counts of gene *k* across cell type *j* for donor *i*. To facilitate the biological interpretability of factors and reduce noise in the tensor formed, we removed genes that are either not provided with HGNC symbols or not labeled as protein-coding in the HGNC database^66^. We also filtered out genes with a total count of less than 50 across all cells. The resulting tensor was total-count normalized, so each sample-cell type pair has 10 ^6^ counts. We then apply C-ZIPTF on the tensor formed.

Briefly, C-ZIPTF employs a Bayesian approach to factorize tensor χ into a set of factors corresponding to donor, cell type, and gene modes by modeling the multidimensional count data using a zero-inflated Poisson distribution. Pseudo-bulk counts χ are modeled as draws from a Zero-inflated Poisson distribution, i.e., for every *ijk*, 1 ≤ *i* ≤ *N*_*D*_, 1 ≤ *j* ≤ *N*_*C*_, 1 ≤ *k* ≤ *N*_*G*_ :

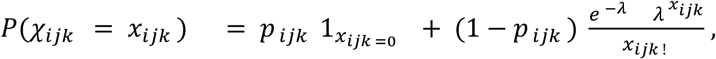

where *x*_*ijk*_ ≥ 0, *λ*_*i*j*k*_ is the expected Poisson count, and *p*_*ijk*_ is the probability of extra zeros. As an abbreviation, it is written as χ_*i*j*k*_ ≈ *ZIP* (*λ*_*ijk*_, *p*_*ijk*_). C-ZIPTF was applied to the pseudo-bulk tensor with a rank of 24. The factorization achieved a high explained variance of 0.916 and a strong cophenetic correlation of 0.931.

The gene latent factors *g*^*r*^, 1 ≤ r ≤ R, represent ranked lists of genes with associated weights. They reveal the expression pattern by capturing co-regulation and co-expression patterns among groups of genes. To identify the top genes defining each factor, we first normalized the gene mode factor matrix such that the L1 norm of each factor equaled 1. We then filtered out genes based on two criteria: (1) genes whose maximum loading across all factors was less than the median of all gene loadings and (2) genes whose entropy was less than the median entropy plus twice the median absolute deviation. Additionally, we conducted gene set enrichment analysis of these factors using GSEApy^67^ in Python, offering a broader perspective on the associated biological functions and pathways. We assess the loadings in *d*^*r*^ to determine the activity level of the GEP in individual donors. Similarly, the cell type loadings in *c*^*r*^ indicate the activity level of the GEP across different cell types.

#### Intratumor heterogeneity analysis

To define the core gene expression programs in HRS cells, we conduct matrix factorization using Zero-Inflated Poisson Tensor Factorization. We subset the data to include only HRS cells and obtain a two-dimensional matrix of cells × genes. Here again, we remove genes that are either not provided with HGNC symbols or not labeled as protein-coding in the HGNC database^66^ and filter out genes with a total count of less than 50 across all cells. This matrix was then total-count normalized so that each cell had 10^6^ counts. We then apply ZIPTF to this matrix, as matrices are essentially two-way tensors. To identify the top genes for each factor, we normalize the gene mode factor matrix such that the L1 norm of each factor equals 1. We then filter out genes based on two criteria: genes whose maximum loading across all factors is less than the median of all gene loadings and genes whose entropy is less than the median entropy plus twice the median absolute deviation. The programs are then annotated manually using gene set enrichment analysis and prior knowledge.

#### Ligand receptor analysis

Ligand receptor interaction inference was performed using the LIANA^68^ tool on the single nucleus RNAseq dataset using the OmniPath^69^ Consensus ligand-receptor interaction database at intermediate cell type annotation hierarchy (“cell_types_level_3”). Ligands or receptors expressed in less than 5% of cells per cluster were excluded. For multimeric receptor complexes, the expression of individual subunits was considered, i.e., receptor complex annotations were not used to reduce the stringency of this analysis due to the sparsity of this annotation and expression levels of receptor complex members. Interactions were prioritized using expression (SingleCellSignalR LRscore^70^, > 0.4), cell type specificity (NATMI edge specificity^71^, > 0.01), and ligand-receptor colocalization metrics (described later). LRscore is a non-negative regularized score comparable between datasets (ranges from 0-1) and reflects the expression magnitude of ligand and receptor. The NATMI edge specificity score ranges from 0 to 1, where 1 means the ligand and receptor are uniquely expressed in each pair of cell types.

### Spatial transcriptomics data analysis

#### Slide-seqV2 alignment and quality control

Demultiplexing, genome alignment, and spatial barcode matching were performed using Slide-seq tools pipeline^53^. The resulting count and spatial barcode matrices were loaded into Seurat for downstream analysis. The “stray” barcodes not present in the densely packed center of the array were cropped out and excluded from analysis by manual inspection of each array.

#### Visium data processing

The demultiplexed FASTQ files were aligned against GRCh38-2020-A human transcriptome reference, and expression count matrices were generated using Space Ranger (v1.3.1). Space Ranger was also used to align the acquired images with the spatially resolved sequencing data.

#### Cell type deconvolution

RCTD v2.0.0, part of the spacexr v2.0.2 package ^72,73^ was used for cell type deconvolution. For the slide-seqV2 dataset, RCTD was run in doublet mode for each sample individually (replicate mode) with the appropriate snRNAseq dataset (lymph node or Hodgkin lymphoma) with corresponding broad cell type labels (level 3) that were assigned as previously described. Some specific cell types were grouped into more coarse cell type classes to allow the model to fall back to the cell class if classification at the level of the granular cell type is not possible. CD4+ and CD8+ T cells were grouped under the T cell class, fibroblasts and follicular dendritic cells were grouped under the stromal cell class, monocytes/macrophages and myeloid dendritic cells were grouped under the “myeloid cell” class, and the lymphatic and vascular endothelial cells were grouped under the “vascular” cell type class. For the Visium dataset, RCTD was run in full mode.

#### CosMx image processing and feature extraction

Raw image processing and feature extraction were performed using the SMI data processing pipeline ^54^, which includes registration, feature detection, and localization. 3D rigid image registration was made using fiducials embedded in the samples matched with the fixed image reference established at the beginning of the SMI run to correct for any shift. Secondly, the RNA image analysis algorithm was used to identify reporter signature locations in the X, Y, and Z axes along with the assigned confidence. The reporter signature locations and the associated features were collated into a single list. Lastly, the XYZ location information of individual target transcript was extracted and recorded in a table by secondary analysis algorithm, as previously described^54^.

#### CosMx cell segmentation

The Z-stack images of immunostaining + DAPI were used to draw cell boundaries on the samples. A cell segmentation pipeline using the Cellpose machine learning algorithm^74,75^ was used to accurately assign transcripts to cell locations and subcellular compartments. The transcript profile of individual cells was generated by combining target transcript location and cell segmentation boundaries. Cells with fewer than 20 total transcripts assigned were omitted from the analysis.

#### Spatial clustering analysis

To determine whether the tumor cells exhibited a random, dispersed, or clustered distribution, we computed Ripley’s statistics using the Ripley’s L function (variance-normalized version of the Ripley’s K statistic) as implemented in the Squidpy package^76^. Arrays with insufficient numbers of tumor cells were excluded from this analysis. The Ripley’s L function was computed for each array with sufficient numbers of tumor cells (>100). The results were visualized and interpreted to assess the spatial organization of tumor cells. For interpretation, we use confidence envelopes: if Ripley’s L at distance d, L(d), exceeds the upper confidence envelope, it indicates statistically significant clustering at that distance; if L(d) falls between the upper and lower confidence envelopes, the distribution is not significantly different from random; and if L(d) falls below the lower confidence envelope, it suggests statistically significant dispersion or regular spacing at that distance.

#### Spatial cell type enrichment

We explore the significant enrichment of cell types surrounding the HRS cells using spatial transcriptomics data, including techniques such as Slide-seq and the CosMx Spatial Molecular Imager (SMI).

We construct a neighborhood graph by considering the spatial locations of the beads (voxels). We count the number of beads classified as cell type *i* within *r* ≥ 0 microns to HRS cells, denote it as *N*_*r*_^0^(*C*_*i*_). To evaluate the significance of enrichment and depletion of a given cell type *i*, we randomly permute the cluster labels while maintaining the connectivity. We repeat this process and record the total count of cell type *i* within a radius of *r* microns to HRS cells at iteration *j* as *N*_*r*_^j^(*C*_*i*_). We estimate the mean and the variance (*μ*_*r*_ (*C*_*i*_), *σ*_*r*_(*C*_*i*_)) of the counts and calculate the enrichment score as following

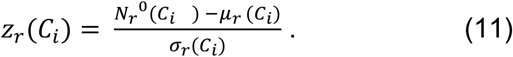

We also perform a one-tailed p-value test to assess the enrichment (or depletion) of cell types. Let *H*(.) be the indicator function such that *H*(*x*) = 1 if *x* > 0, and 0 otherwise, then we

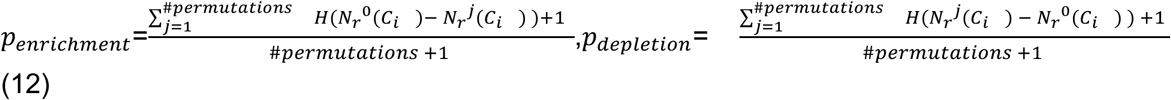

We individually perform this test for each tissue and calculate the median of the enrichment scores across all the tissues. The resulting p-values from each tissue are combined using Fisher’s method (Fisher’s combined probability test) by utilizing the scipy.stats.combine_pvalues function from the SciPy library. To control the expected proportion of false discoveries among the rejected hypotheses, we apply the Benjamini-Hochberg Procedure for False Discovery Rate (FDR) correction. This adjustment ensures a more accurate control of the overall false positive rate in our analysis.

#### Cell-type specific spatial differential expression

After cell type deconvolution using RCTD, we performed cell type specific differential expression analysis across the entire multi-sample multi-replicate slideseqV2 dataset using C-SIDE^73^. The samples represent tumors from different patients, and replicates refer to other sections/arrays from the same tumor. We hypothesized that the local density and proximity of tumor cells are plausible predictors of gene expression in non-tumor cells. Therefore, we first calculated the normalized (min-max normalization per array) density of HRS cells for each pixel within a 200 µm radius. Second, for each pixel, we calculated the inverse of the distance to the nearest tumor cell (if distance = 0, proximity = 1). We then ran CSIDE in replicate mode (run.CSIDE.replicates) using these two orthogonal continuous explanatory variables to test for differentially expressed genes in non-tumor cells as a function of local proximity or density of tumor cells. Default parameters were used except for the normalized expression threshold for gene selection (gene_threshold = 1e-06) and the minimum number of cells of each cell type to be used (cell_type_threshold = 0). As implemented in CSIDE (CSIDE.population.inference), population-level differential expression analysis was performed across all samples to identify genes that consistently show a similar pattern of differential expression across multiple samples. Default parameters were used except the minimum number of groups (if use.groups) for which a gene must converge (MIN.CONV.GROUPS = 3). Genes with a log2 fold change value of > 0.5 and a q value of < 0.05 were considered significantly differentially expressed. Cell type-specific genes were prioritized to minimize the potential of false positive genes that may result from cell type misclassification and/or diffusion. Cell-type specific genes were enriched by setting the ct_prop (minimum ratio of expression in a given cell type compared to all other cell types) threshold at a stringently high level (ct_prop > 0.5).

#### Ligand receptor colocalization

To test the significance of colocalization of ligand-receptor pairs as a measure of interaction between cell types expressing those, we set up a series of permutation tests. We used count matrices of slide-seq libraries from lymphoma samples and ligand-receptor interacting pairs inferred from single-nuclei data using LIANA as our starting point. To remove potential bias due to empty regions of the arrays, which contain very few counts, we removed beads with the maximum number of detected genes below a certain threshold. This threshold was determined by finding the maximum between 150 (heuristically determined to reflect empty tissue sections in most arrays) or the value of the lower quartile of gene counts per bead (a higher value on a few arrays that had unusually high capture of transcripts). Furthermore, for each array, we did not analyze pairs where either ligand or receptor had less than 20 representative beads on that array. For specific highly expressed ligand or receptor genes such as B2M or CD3E, we observed that most of the beads on the arrays had nonzero expression. To binarize the expression of such genes into positive and negative categories, we considered beads that had expression below the 75th percentile of counts for that gene as negative. This value was selected heuristically so that only a few highly expressed genes would be affected. This thresholding mechanism would not affect the vast majority of genes that have mostly either counts of 1 or 0 in all beads. Subsequently, we counted the number of receptor-positive beads in an area around each ligand-positive bead as the observed value. We selected 1000 random sets of points equal to the number of ligand-positive beads from the set of beads deemed to contain tissue and counted the receptor-positive beads around each random ligand position. We calculated the significance of this permutation test as (1+n)/(N+1), where n is the number of times the random permutation yielded a higher count than the observed value. We also calculated an effect size equal to the ratio of observed value and mean of values obtained from random permutations. We repeated this procedure across all arrays and over multiple radii. We combined the p-values for each radius using Fisher’s method and corrected them across the entire set of ligand-receptor pairs using the Benjamini-Hochberg method. The combined effect size for each pair and radius was calculated as the average of this value across all arrays. The product of the p-value and the effect size were used as the combined spatial association metric to reflect consistency (across replicates) and magnitude of colocalization (shown in Fig. 3).

## Extended data figures

**Fig. 1:**
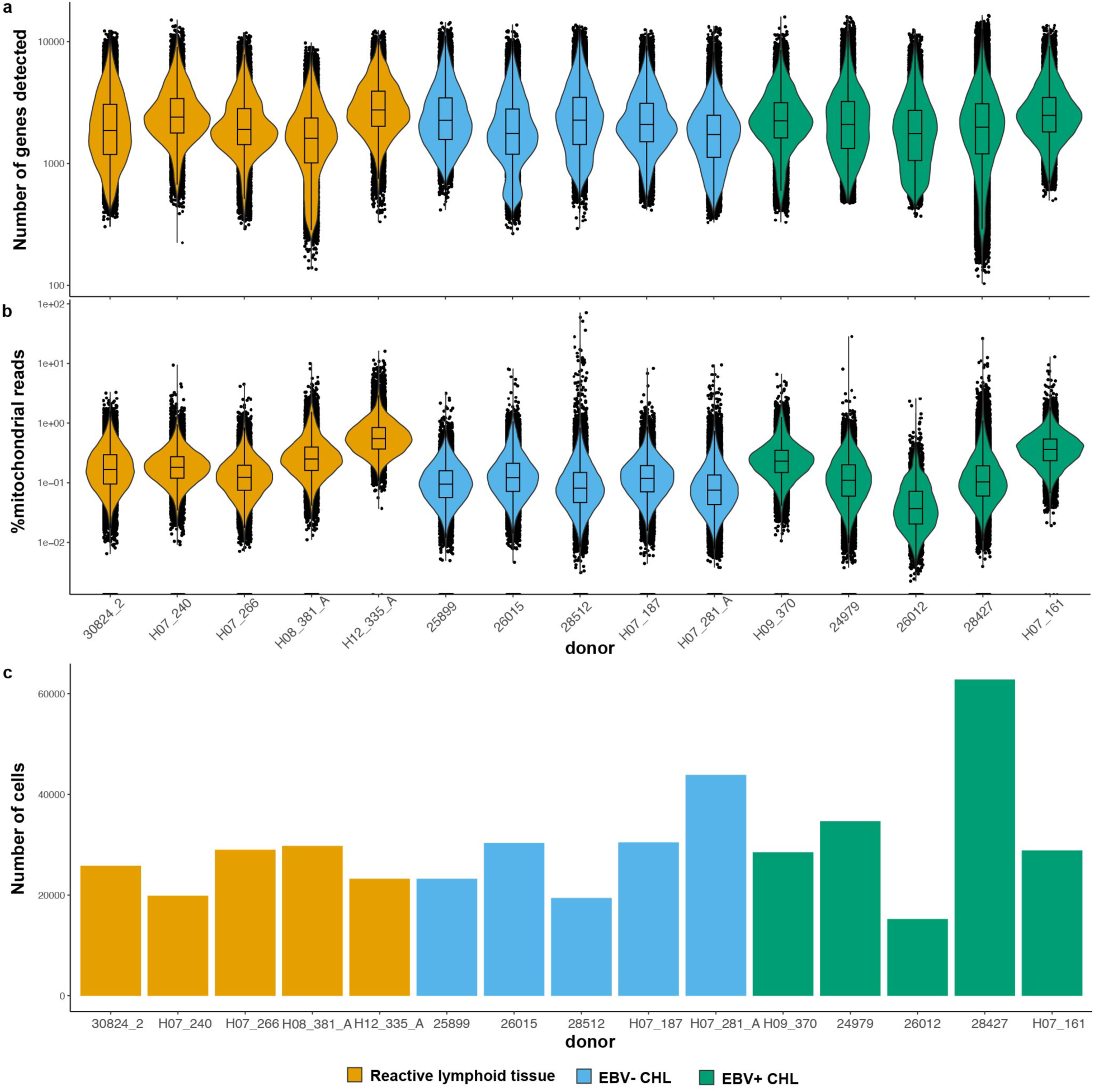
Summary and quality control metrics of snRNAseq dataset. **a**, Violin plots of a, number of genes detected per cell, **b**, % mitochondrial reads, and **c**, barplot of the number of cells passing quality control metrics. Samples are colored based on EBV status (positive or negative) and sample type (reactive lymph node or Hodgkin lymphoma).

**Fig. 2:**
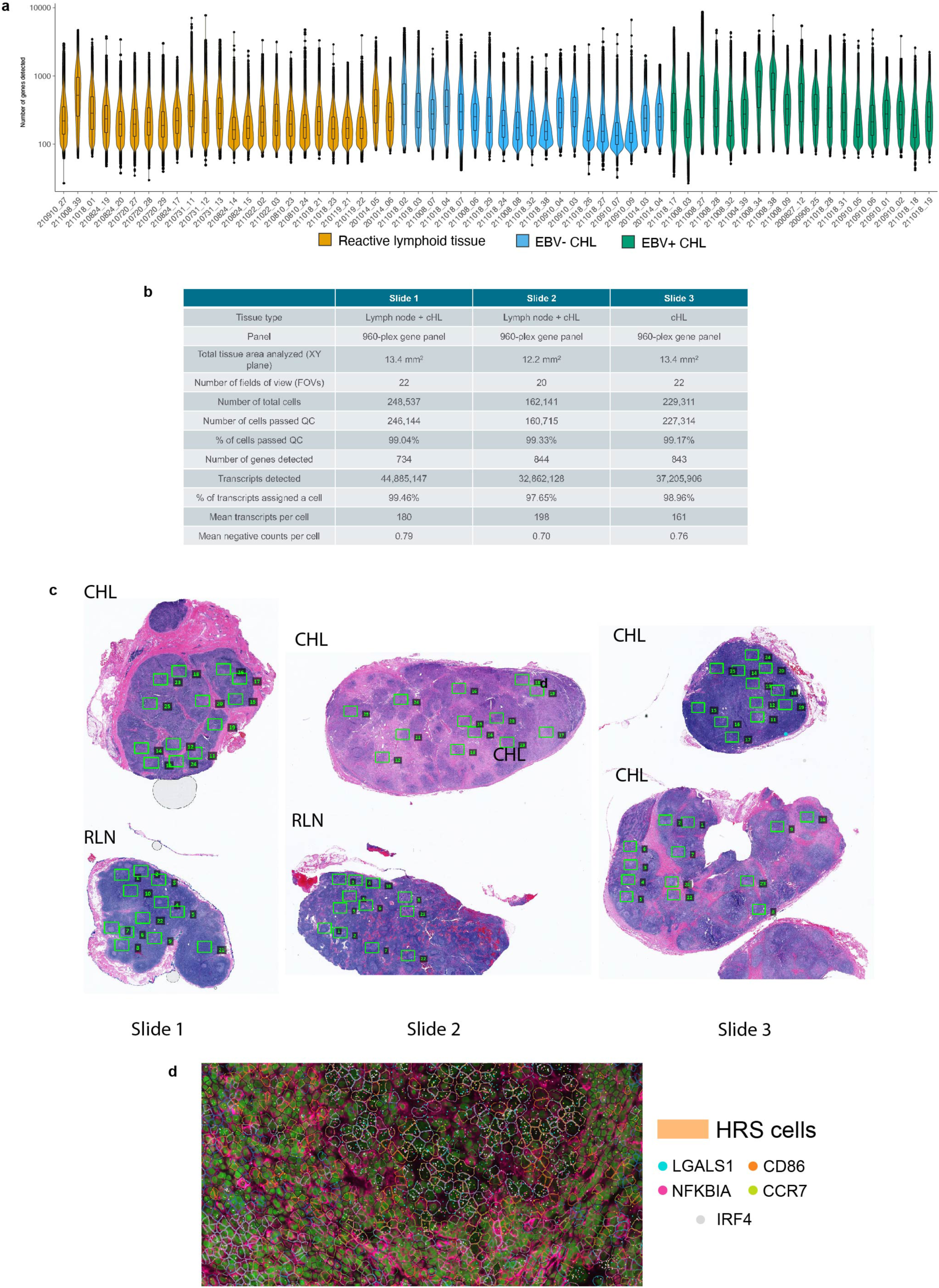
Summary and quality control metrics of the spatial dataset. **a**, Violin plot of the number of genes detected per pixel across all Slide-seqV2 arrays. Samples are colored based on EBV status (positive or negative) and sample type (reactive lymph node or Hodgkin lymphoma). **b**, quality control metrics of the CosMx dataset on a per-slide basis. **c**, Whole slide images of hemotoxylin and eosin stained sections showing the fields of view (green boxes) selected for analysis. **d**, representative field of view (FOV) of a cHL specimen with segmentation masks colored by cell type annotation and spots representative single transcripts colored by gene (HRS cell marker genes). The cells classified as HRS cells are large, multinucleated, and show expression of known marker genes.

**Fig. 3:**
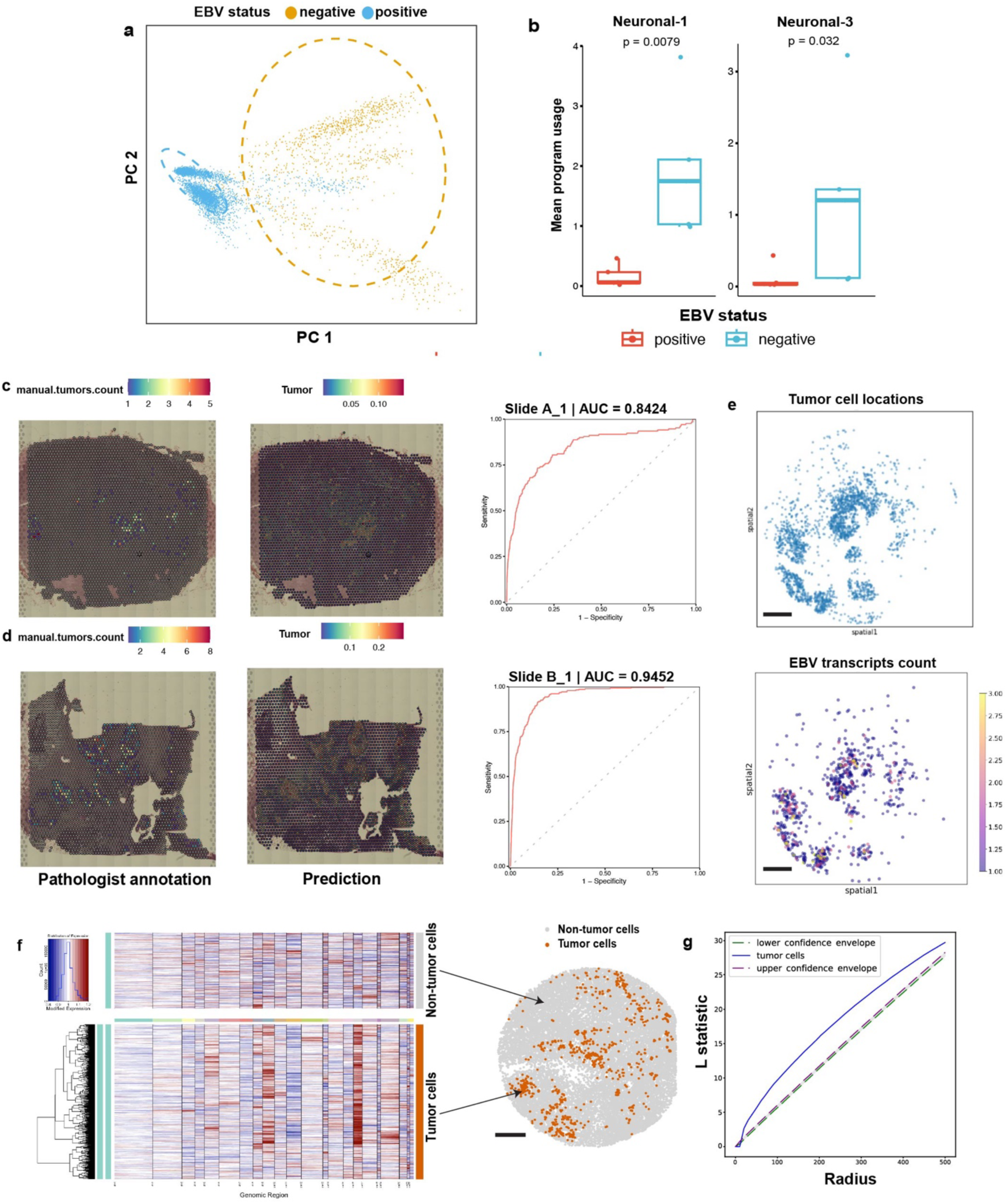
Expression programs and spatial mapping of HRS cells. **a**, Principal component analysis (PCA) plot of HRS cells gene expression profiles colored by EBV status. **b**, boxplot showing mean program usage of the neuroglial programs per donor split by EBV status showing the association between expression of these neuroglial signatures and EBV status. **c, d**, spatial plot of tumor cells determined by morphology by a pathologist (left) and gene expression (RCTD derived tumor cell weights, middle) and area under the receiver operating characteristics curve plot on a Visium spatial gene expression dataset of two Hodgkin lymphoma tumors. **e**, spatial plot of tumor cells based on the expression of known marker genes (top) in an EBV+ Hodgkin lymphoma and spatial plot of EBV-specific transcripts on the same array (bottom). **f**, inferCNV analysis of spatially mapped putative tumor cells (orange spots) with mapped non-tumor cells set as reference cells on a slide-seqV2 dataset of Hodgkin lymphoma showing evidence of copy number variants, including gain of 9p. **g**, Ripley’s L statistic for tumor cells across multiple radii on the array shown in the main fig. 1g indicating evidence of clustering (representative array from donor H12-200). All scale bars: 500 µm.

**Fig. 4:**
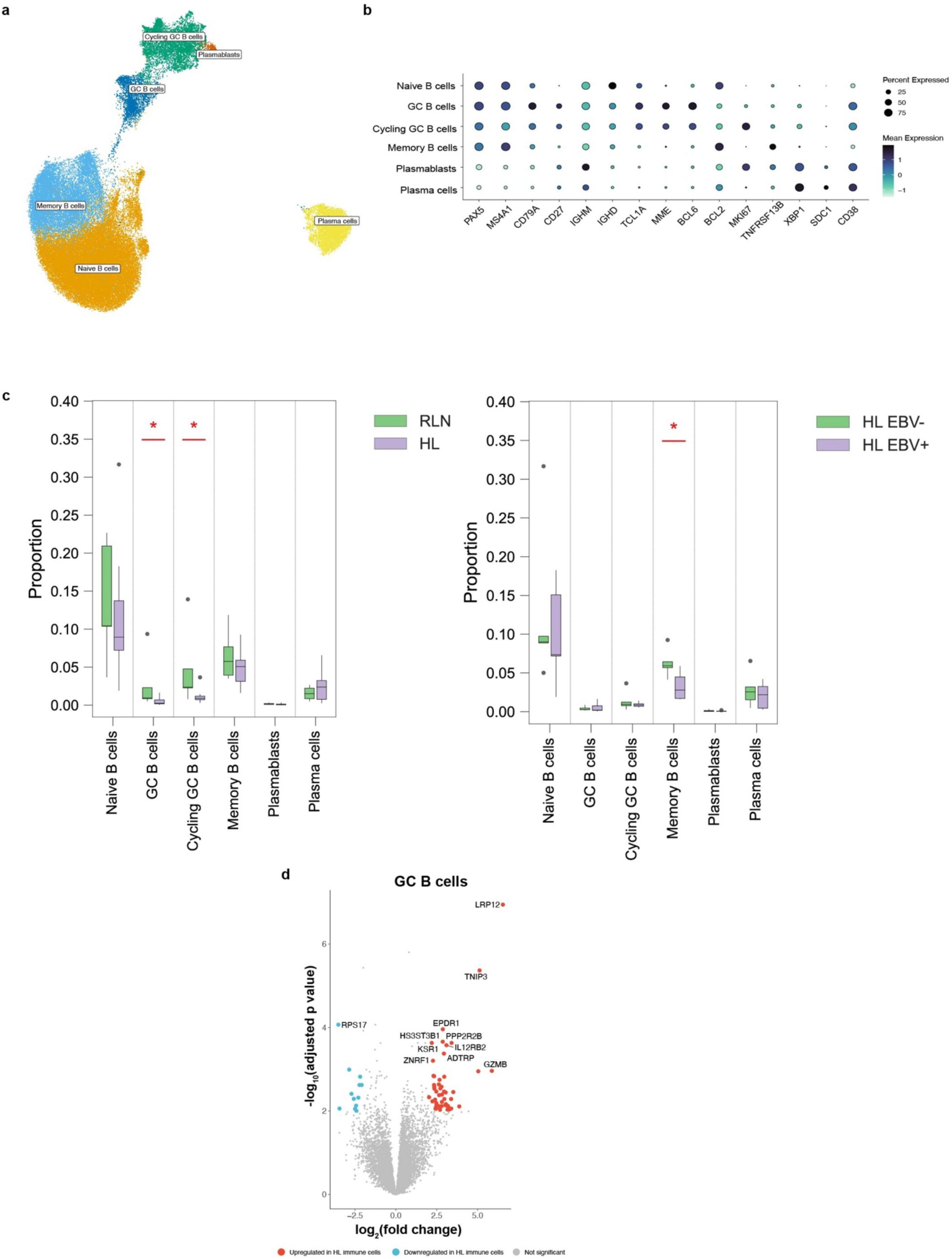
Annotation and compositional analysis of B cells. **a**, UMAP embedding of B cells colored by cell type annotation. **b**, bubble plot showing curated marker genes across all annotated cell type clusters. **c**, compositional analysis of B cell subsets across Hodgkin lymphoma and reactive lymph node samples (left) and EBV positive and negative samples (right). **d**, volcano plot showing differentially expressed genes in B cells in Hodgkin lymphoma samples versus reactive lymph node samples.

**Fig. 5:**
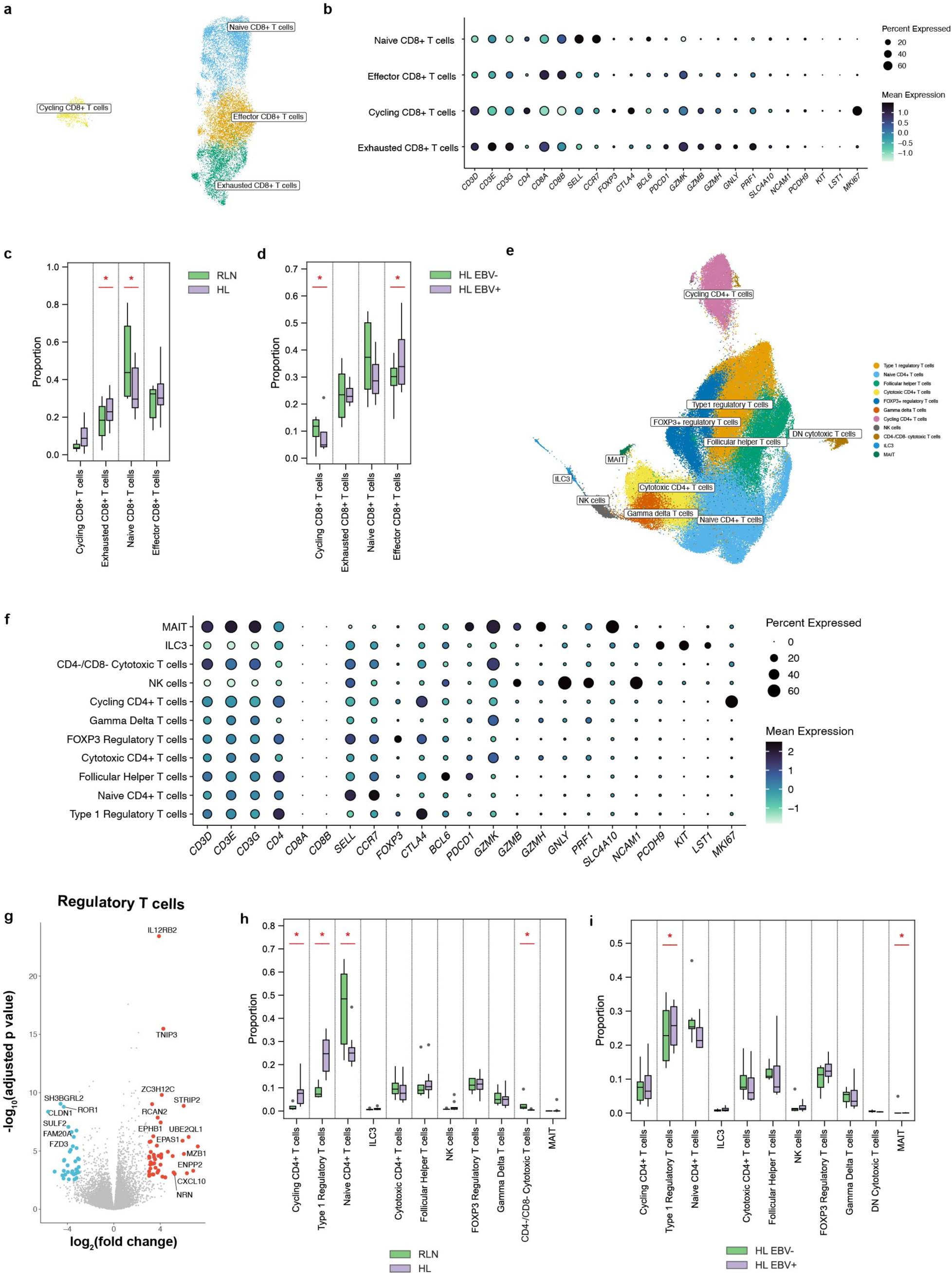
Annotation and compositional analysis of T cells. **a**, UMAP embedding of CD8+ T cells colored by cell type annotation. **b**, bubble plot showing expression of curated marker genes across all annotated cell type clusters in CD8+ T cells. **c, d**, compositional analysis of CD8+ T cell subsets across Hodgkin lymphoma and reactive lymph node samples (c) and EBV positive and negative samples (d). **e**, UMAP embedding of non-CD8+ T cells colored by cell type annotation. **f**, bubble plot showing curated marker genes across all annotated cell type clusters in non-CD8+ T cells. **g**, volcano plot showing differentially expressed genes in regulatory T cells in Hodgkin lymphoma samples versus reactive lymph node samples. **h, i**, compositional analysis of non-CD8+ T cell subsets across Hodgkin lymphoma and reactive lymph node samples (h) and EBV positive and negative samples (i).

**Fig. 6:**
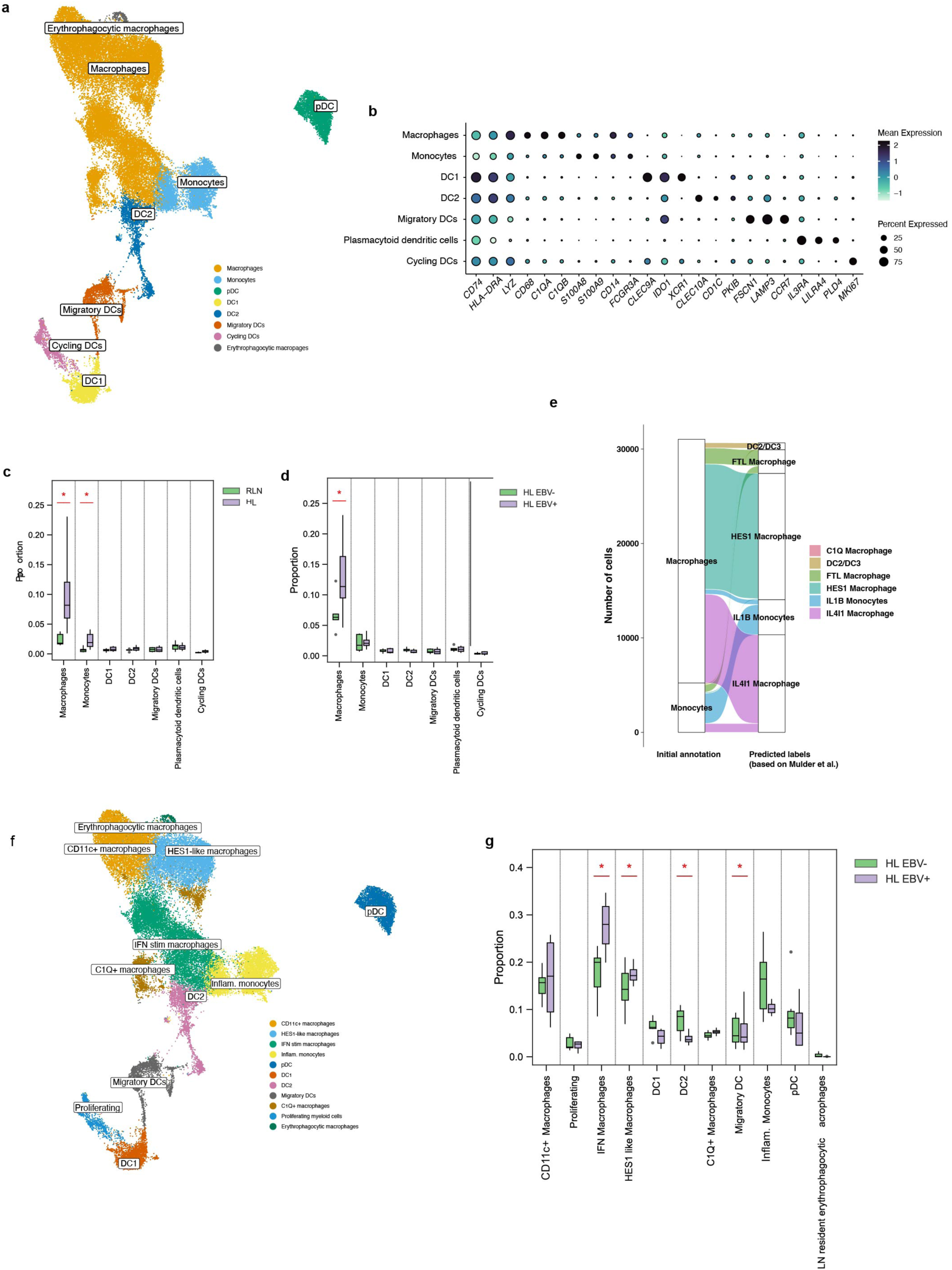
Annotation and compositional analysis of myeloid cells. **a**, UMAP embedding of myeloid cells colored by cell type annotation. **b**, bubble plot showing expression of curated marker genes across all annotated myeloid cell type clusters. **c, d**, compositional analysis of myeloid cell subsets across Hodgkin lymphoma and reactive lymph node samples (c) and EBV positive and negative samples (d). **e**, Sankey plot showing classification of monocytes/macrophages into finer cell types based on label transfer analysis from the MoMac-verse reference dataset^34^. **f**, UMAP embedding with cells colored by finer myeloid cell type annotations based on label transfer analysis from MoMac-verse reference dataset^34^. **g**, compositional analysis based on EBV status on finer myeloid cell type annotations.

**Fig. 7:**
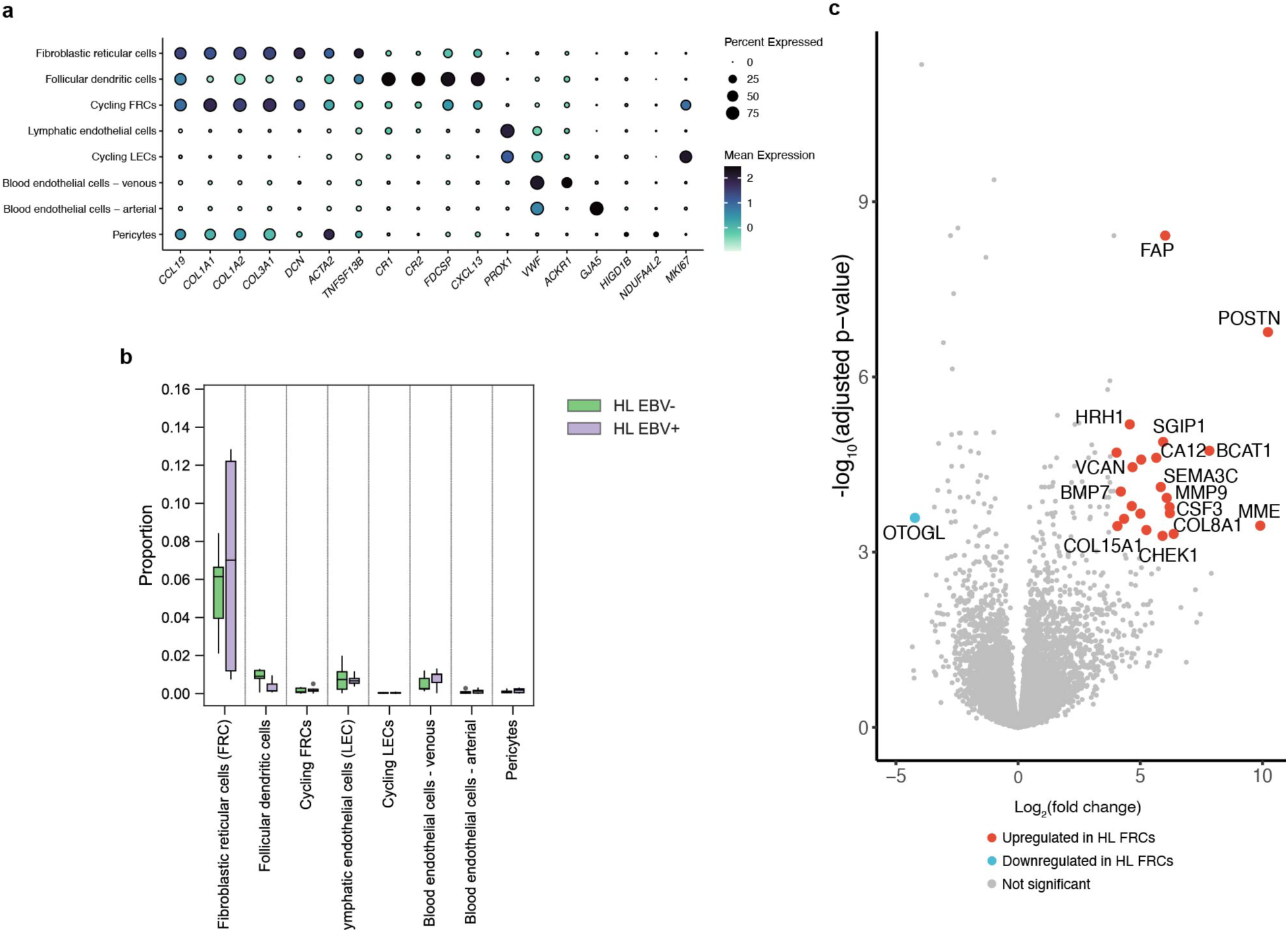
Annotation, compositional, and differential expression analysis of stromal cells. **a**, bubble plot showing expression of curated marker genes across all annotation stromal cell type clusters. **b**, compositional analysis of stromal cell subtypes across EBV positive and negative samples. **c**, volcano plot showing differentially expressed genes in fibroblastic reticular cells in Hodgkin lymphoma versus reactive lymph node samples.

**Fig. 8:**
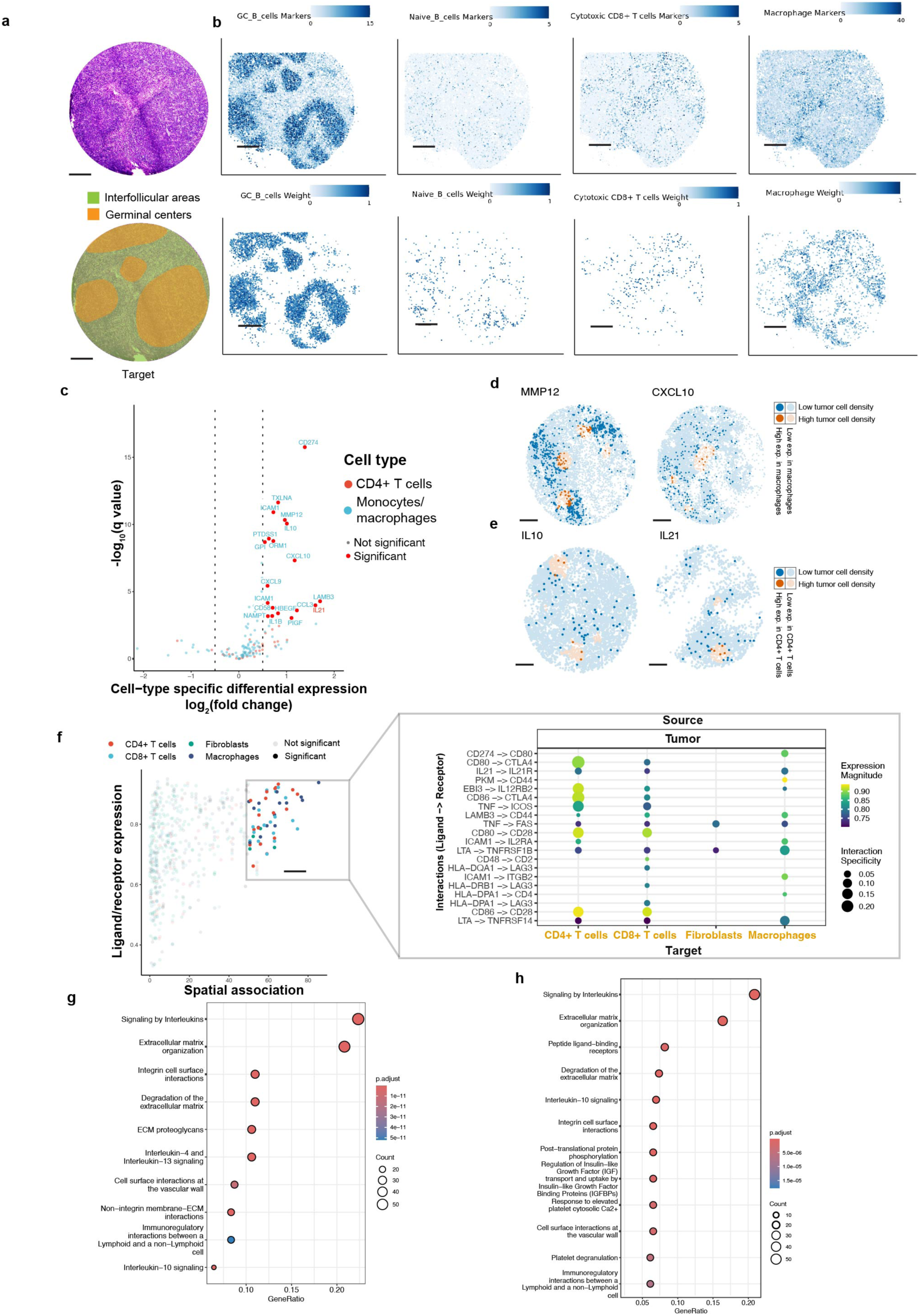
Spatial organization of the immune microenvironment and ligand-receptor interactions around tumor cells. **a**, H&E-stained serial section of a reactive lymph node specimen (top) with histologic annotation of follicular and interfollicular zones (bottom). **b**, spatial plot of marker genes (top row) of various cell types and predicted spatial locations by RCTD (cell type weights; bottom row); same array shown in a. **c**, volcano plot showing cell-type specific differentially expressed genes in relation to HRS cell proximity. A positive log_2_(fold change) indicates genes that show higher expression in HRS cell proximal regions. **d**, representative spatial plots showing expression of MMP12 (left) and CXCL10 (right) in macrophages in tumor cell dense and sparse regions. **e**, spatial plots showing expression of IL10 (left) and IL21 (right) in CD4+ T cells in tumor cell dense and sparse regions. **f**, scatterplot of tumor-to-immune ligand-receptor interactions based on spatial proximity (spatial association metric, see methods) and normalized ligand/receptor expression (sca LRscore metric). Interactions that show high ligand/receptor expression and spatial proximity are called out and shown on the right as a bubble plot. **g, h**, enrichment analysis of spatially associated ligand-receptor interactions (g) and genes upregulated in relation to HRS cell density/proximity (g). All scale bars: 500 µm.

**Fig. 9:**
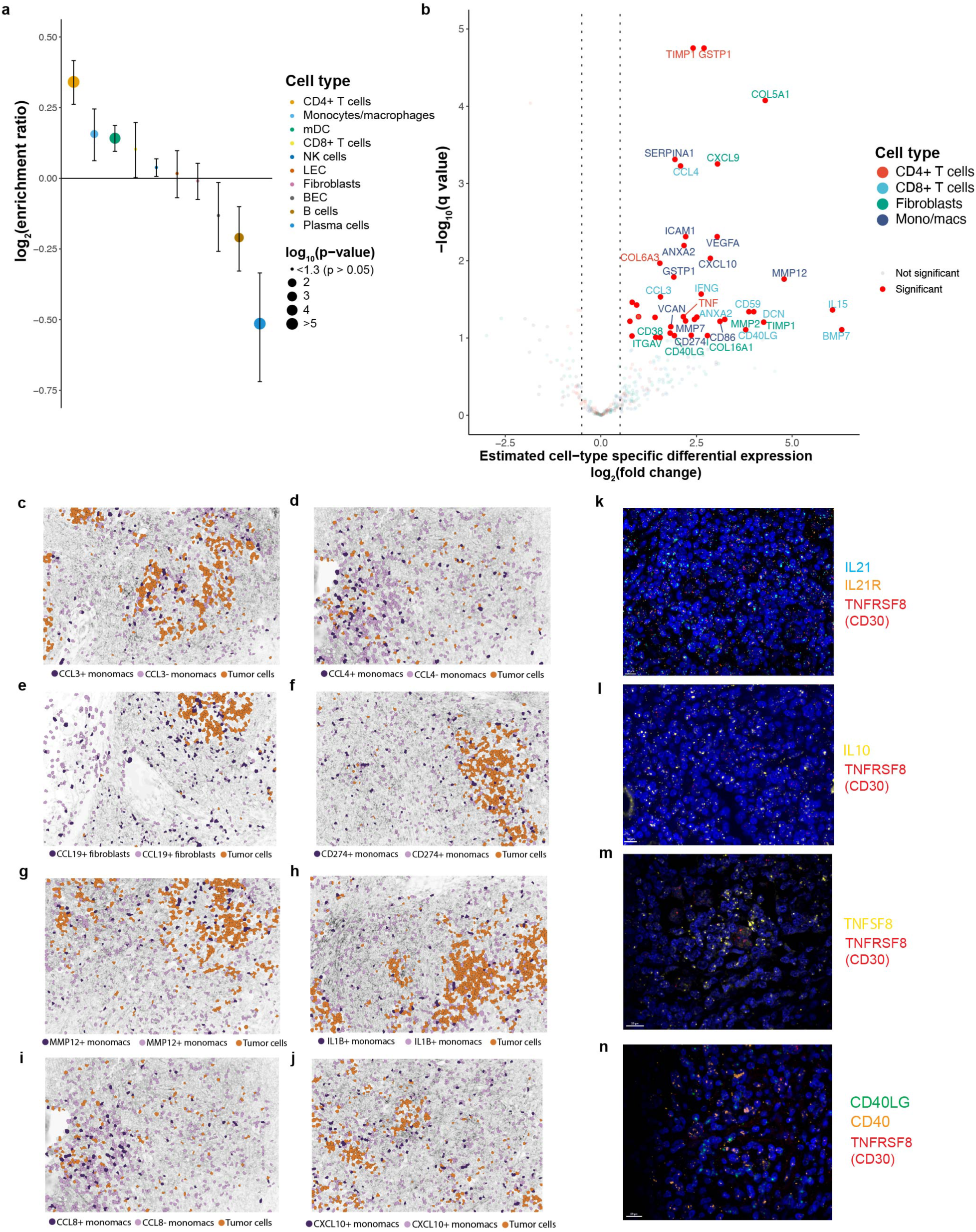
Validation of spatially associated cell types and ligand-receptor interactions using imaging transcriptomics. **a**, Enrichment ratios of immune cell types (colored by cell type) aggregated across the entire multi-sample, multi-replicate cosMx dataset. The size of the points indicates the P value (one sample t-test, two-tailed); mean ± s.e.m. is depicted. **b**, volcano plot showing cell-type specific differentially expressed genes in relation to HRS cell proximal regions in the CosMx dataset. A positive log_2_(fold change) indicates higher expression in HRS cell proximal regions, and points are colored by cell type. **c-j**, representative fields of view from the CosMx dataset showing enrichment of the indicated ligands in the relevant cell types around tumor cells. The data are representative of four independent experiments on four different patients. **k-n**, RNA in-situ hybridization analysis showing spatial association of microenvironment-derived ligands (IL21, CD40LG, TNFSF8, IL10) and the cognate receptors (IL21R, CD40) expressed on HRS cells. The data are representative of three independent experiments on at least two different patient samples. All scale bars: 10 µm.

**Fig. 10:**
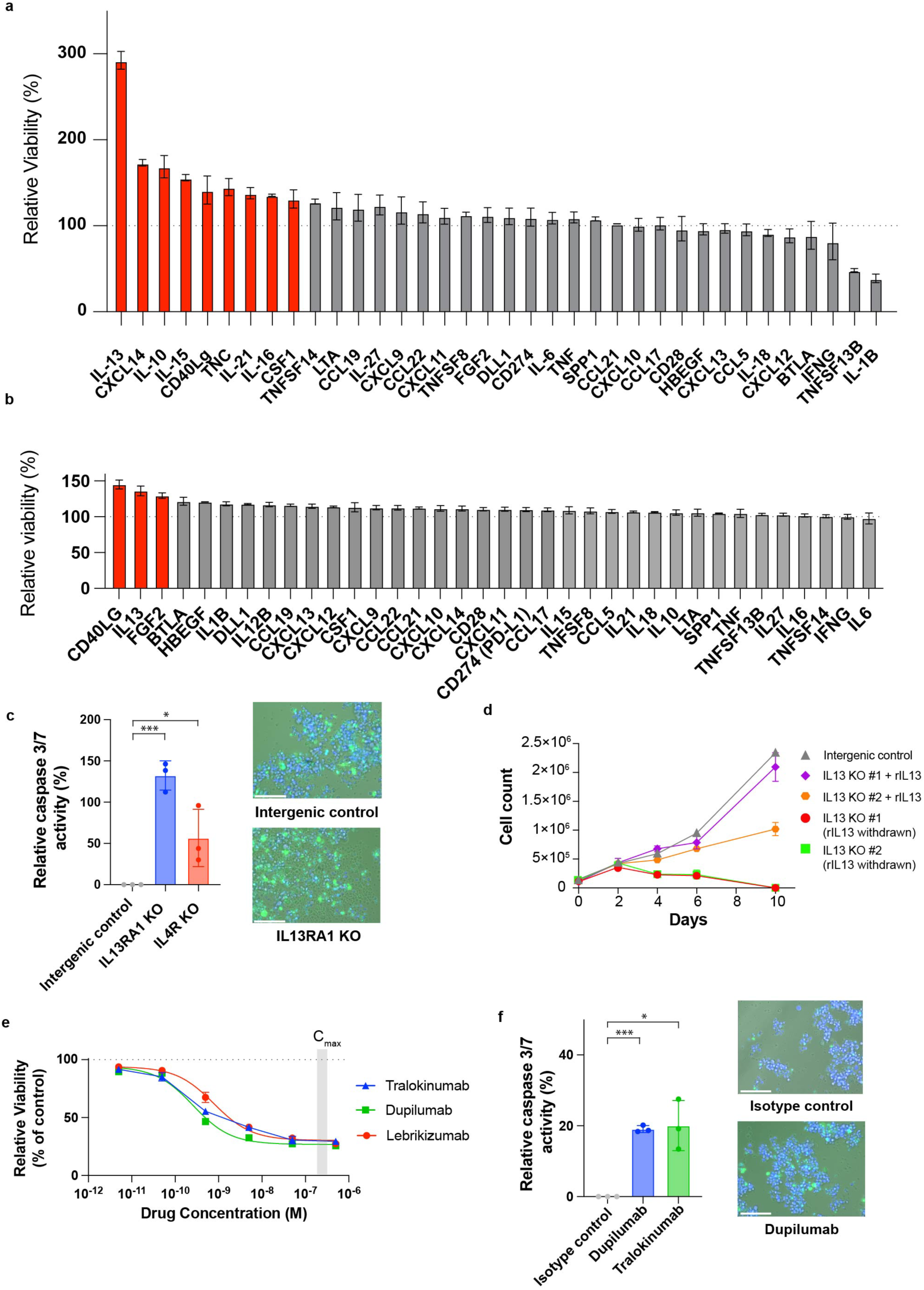
Experimental validation of predicted microenvironment-derived growth factors. **a**, Barplot showing the effect of recombinant factors on the relative viability (normalized to untreated control) of L1236 cells at three days at low-density growth conditions. **b**, barplot showing the effect of recombinant factors on the relative viability (normalized to untreated control) of UHO1 cells at three days grown under low serum conditions; for a and b, mean ± s.d. is depicted (n = 3). *P* < 0.05 is considered significant; two-way analysis of variance (ANOVA) followed by Dunnett’s multiple comparison test. **c**, mean nuclear CellEvent Caspase 3/7 green fluorescence in IL4R and IL13RA1 knockout L1236 cells relative to intergenic control (24 hours after transduction/selection, n = 3) and representative composite images of the intergenic control and IL13RA1 knockout cells (channels: blue - DAPI, green – CellEvent green, transmitted light). **d**, the effect of withdrawing recombinant IL13 from IL13 knockout L1236 cells that were cultured in the presence of recombinant IL13 for 8 days (from the experiment shown in main fig. 4g); mean ± s.d. is depicted (n = 2). **e**, dose-response analysis of tralokinumab, dupilumab, and lebrikizumab on UHO1 HRS cells (n = 4); mean ± s.e.m. is depicted. The C_max_ range of these antibodies reported in clinical trials is indicated in grey. **f**, mean nuclear CellEvent Caspase 3/7 green fluorescence of L1236 cells treated with 0.5 µM Dupilumab or Tralokinumab relative to isotype control antibody (24 hours after treatment, n = 3) and representative composite images of the isotype control and dupilumab treated cells (channels: blue - DAPI, green – CellEvent green, transmitted light); For c and f, mean ± s.d. is depicted, the data are representative of two independent experiments and *P* values are determined by two-tailed t-tests. \**P* < 0.05, \*\**P* < 0.01, \*\*\**P* < 0.001. All scale bars: 150 µm.

## Data and code availability

All single-cell and spatial transcriptomics datasets will be deposited on the Broad Institute Single cell Portal at the time of publication. The raw sequencing data will be available through a controlled access data repository (dbGAP). Expression count matrices will be deposited in GEO. Code to reproduce all the analyses in this manuscript is available on Github: https://github.com/klarman-cell-observatory/chl_analysis.

## Acknowledgments

This work was supported by funding from the Damon Runyon Cancer Research Foundation (PST: 46-24), the Lymphoma Research Foundation (V.S.), the Eric and Wendy Schmidt Center at the Broad Institute of MIT and Harvard (N.T.), and the National Cancer Institute R35CA242457 (T.R.G.). V.S. also thanks Dr. Jon Aster and Dr. Kathleen Burns for their mentorship and the Brigham and Women’s Hospital Department of Pathology and the Dana-Farber Cancer Institute Department of Oncologic Pathology for generously supporting his research training.

## Author contributions

V.S., F.C., and T.R.G. conceived the study with input from A.L, M.A.S, and S.J.R. V.S. performed experiments with help from S.S., S.H., I.B., J.A.W., H.M., N.N. and G. N. V.S., D.C. and N.T. performed analyses with help from M.B., C.U., D.C., J.A.W. and O.A. G.P. performed key immunohistochemical studies. M.A.S. and A.L. contributed key samples. V.S. and T.R.G. wrote the paper with contributions from all the authors.

